# Inactivation of mitochondrial complex IV in *Physcomitrium patens* reveals the essential role of respiration in coordinating plants metabolism

**DOI:** 10.1101/2024.05.06.592826

**Authors:** Antoni M. Vera-Vives, Marco Mellon, Libero Gurrieri, Philipp Westhoff, Anna Segalla, Edoardo Bizzotto, Stefano Campanaro, Francesca Sparla, Andreas P.M. Weber, Alessandro Alboresi, Tomas Morosinotto

**Affiliations:** Department of Biology, University of Padova, Italy; Department of Pharmacy and Biotechnology (FABIT), University of Bologna, Italy; Plant Metabolism and Metabolomics Laboratory, Cluster of Excellence on Plant Science (CEPLAS), Heinrich Heine University, D-40225 Düsseldorf, Germany; Institute of Plant Biochemistry, Cluster of Excellence on Plant Science (CEPLAS), Heinrich Heine University, Düsseldorf, Germany

## Abstract

Photosynthetic organisms use sunlight as energy source but rely on respiration during the night and in non-photosynthetic tissues. Respiration is also active in photosynthetically active cells, where its role is still unclear due to a lack of viable mutants.

Plants lacking cytochrome c oxidase (complex IV) activity are generally lethal but were here isolated exploiting the possibility of generating knockout lines through vegetative propagation in the moss *Physcomitrium patens*.

The mutants showed severely impaired growth, with an altered composition of the respiratory apparatus and increased electron transfer through the alternative oxidase. The light phase of photosynthesis remained largely unaffected while the efficiency of carbon fixation was moderately reduced. Transcriptomic and metabolomic analyses showed that the disruption of the cytochrome pathway had broad consequences for carbon and nitrogen metabolism. A major alteration in nitrogen assimilation was observed with a general reduction in amino acid abundance. A partial rescue of the growth could be obtained by growing the plants with an external supply of amino acids but not with sugars, demonstrating that respiration in plant photosynthetic cells plays an essential role at the interface between carbon and nitrogen metabolism and a key role in providing carbon skeletons for amino acid biosynthesis.

## Introduction

Photosynthetic organisms use light energy as their primary source of energy to synthesize the ATP and the reducing power (NADPH) needed to support their entire metabolism and especially the fixation of CO_2_ into organic molecules. This metabolic reaction is responsible for most of the Earth’s primary production, providing the chemical energy to support most lifeforms. Photosynthetic organisms also rely on respiration to meet energy demands at night (i.e., in the absence of light) or in tissues that are not photosynthetically active like roots or seeds during germination.

Oxidative phosphorylation is the main metabolic pathway responsible for generation of ATP in heterotrophic cells and in eukaryotes it is localized in mitochondria. This process is fed by reducing power originating from the oxidation of organic acids and leads to the release of CO_2_ and to the reduction of O_2_ to water. Oxidative phosphorylation is catalysed by five major large multi-subunit complexes embedded in the inner mitochondrial membrane: the NADH dehydrogenase complex (complex I, CI), the succinate dehydrogenase complex (complex II, CII), the cytochrome c reductase complex (complex III, CIII), the cytochrome c oxidase complex (complex IV, CIV), and the ATP synthase (complex V, CV). Together, CI to CIV form the mitochondrial electron transfer chain (mETC) or respiratory chain, which catalyses the transfer of electrons from NADH or FADH_2_ to molecular oxygen. CI, CIII, and CIV of the respiratory chain are proton translocators and the mETC activity drives the generation of an electrochemical gradient across the inner mitochondrial membrane which is subsequently used by CV to catalyse the phosphorylation of ADP to ATP.

In addition to its role in supporting energy metabolism in heterotrophic tissues or in the dark (Millar et al., 2011), plant respiration has been shown to be active also in the light and to influence photosynthesis by acting as a sink for excess of electrons coming from the chloroplasts thanks to the action of the malate-oxaloacetate valve, which allows the exchange of reducing power between organelles (Alric & Johnson, 2017; Igamberdiev, 2020; Igamberdiev & Bykova, 2023). It has been shown that mitochondrial respiration activity increases under illumination (De Col et al., 2017) and directly affects photosynthetic electron transport (Larosa et al., 2018) and chloroplast ATPase activity (Mellon et al., 2021). In diatoms, a group of marine algae, the energetic coupling between plastids and mitochondria is essential to drive CO_2_ assimilation (Bailleul et al., 2015). A growing body of evidence suggests that respiration plays an important biological role in plants, even under illumination, and that its activity is closely linked to photosynthesis.

Mitochondrial respiration is essential to supply ATP to the cytosol, even in photosynthetically active cells, as recently demonstrated in *P. patens* (Vera-Vives et al., 2024). Indeed, the inner mitochondrial membrane contains an efficient ADP/ATP exchange system (Fricaud et al., 1992; Gout et al., 2014). Although chloroplasts can indirectly export ATP to the cytosol under some conditions (Gardeström & Igamberdiev, 2016; Heber & Santarius, 1970; Stocking & Larson, 1969), mitochondrial respiration is an essential source of ATP for sucrose synthesis also in illuminated, photosynthetic tissues (Gardeström et al., 2002; Kromer, 1995; Noguchi & Yoshida, 2008).

Plant mitochondria have additional roles in stress responses, including abiotic stress tolerance and the orchestration of programmed cell death (Aken, 2018; Scott & Logan, 2008; Suzuki et al., 2012),in cellular redox regulation and the management of reactive oxygen species (ROS) (Geigenberger & Fernie, 2014), and in the biosynthesis of key metabolites such as amino acids, fatty acids, vitamin co-factors or tetrapyrroles (Fernie et al., 2020; Mackenzie & McIntosh, 1999; Palmer, 1979). Accordingly, several solute transporters are present in the inner mitochondrial membrane (Fernie et al., 2020).

The synthesis of amino acids in the cytosol and plastids by the glutamine synthetase/glutamine:2-oxoglutarate aminotransferase (GS/GOGAT) enzymes requires a supply of carbon skeletons, which are obtained through the direct or indirect export of alpha-ketoglutaric acid from the mitochondrial tricarboxylic acid cycle (TCA cycle) (Day et al., 2004; Gauthier et al., 2010). Amino acids can also be used as a source of electrons to fuel the mETC during carbon starvation (Cavalcanti et al., 2017; Szal & Podgórska, 2012; van Aken, 2021).

Despite its biological relevance, the role of mitochondrial respiration in plants under illumination is far from being understood. A major limitation in advancing knowledge in this area has been the lack of viable land plant mutants with depleted respiration. While there is a collection of mutants depleted in one or more respiratory complexes in the green alga *Chlamydomonas reinhardtii*, as reviewed by Salinas et al., 2014, complete knockout mutants in land plants are only available for complex I in *Arabidopsis thaliana* (Fromm, Braun, et al., 2016; Meyer et al., 2009; Pétriacq et al., 2017), *Nicotiana sylvestris* (Brangeon et al., 2000; Gutierres et al., 1997), and the moss *Physcomitrium patens* (Mellon et al., 2021). Full knockout mutants for complexes II-V have never been isolated in seed plants because of the negative effects on embryo development and seed germination.

The lethal consequences of complex IV depletion have been reported several times (Gras et al., 2020; Kolli et al., 2019; Mansilla et al., 2015; Steinebrunner et al., 2011). Only plants with a reduced accumulation of specific subunits of complex IV have been isolated and they exhibited decreased viability, developmental inhibition, sterility, and impaired seed germination (Kolli et al., 2019). These phenotypes are consistent with the energetic depletion of non-photosynthetic tissues. A *cod1* mutant in Arabidopsis has been reported with undetectable levels of complex IV activity (Dahan et al., 2014). However, *cod1* seeds failed to germinate, and plants could only be obtained by *in vitro* embryo rescue (Dahan et al., 2014). Germination occurred after several weeks (approx. 3 months) of incubation, and subsequent development of seedlings was very slow and disordered (Dahan et al., 2014). The strongly altered development of these plants did not allow a full assessment of the metabolic impact of respiration deficiency in photosynthetic tissues.

The lack of viable complex IV-depleted plants was addressed using the moss *Physcomitrium patens* where knockout (KO) mutants can be generated by homologous recombination in vegetatively propagated tissues cultivated under continuous light. Most *P. patens* tissues are haploid, allowing direct assessment of the final phenotype associated with any genetic modification by bypassing sexual reproduction and thus respiration-dependent developmental stages such as fertilization and seed or spore germination. Most of the life cycle of *P. patens* consists of photosynthetically active cells, and even rhizoids still contain some active chloroplasts (Sakakibara et al., 2003), making it a highly suitable model for isolating mutants with potentially altered mitochondrial functions, as recently demonstrated with the isolation of a complex V KO mutant (Vera-Vives et al., 2024).

For generating mutants in *P. patens* depleted in complex IV activity we chose the protein COX11 as a target. COX11 is an assembly factor required for the insertion of two copper ions into the COX1 subunit, which forms the Cu_B_ centre of the complex IV catalytic core (Meyer et al., 2019, Figure 1A). COX11 is well conserved in different organisms with respiratory activity (Esposti, 2020; Timón-Gómez et al., 2018). COX11 activity has been shown to be essential for complex IV functionality, and its depletion led to respiratory null mutants in yeast (Banting & Glerum, 2006), motivating its selection as a target for generating complex IV depleted plants. A complete knockout could not be achieved in Arabidopsis, while the reduction of COX11 levels by knockdown approaches led to defective embryo development (Radin et al., 2015).

Here, we present the generation and characterization of novel *P. patens* plants defective in COX11, which show completely abolished complex IV activity and altered respiratory activity. The mutation caused a strong alteration in carbon metabolism, with an inability to mobilize energy reserves. Nitrogen metabolism was also altered, due to a lack of carbon skeletons for amino acid biosynthesis, demonstrating that mitochondrial metabolic activity is essential even in photosynthetically active cells.

## Results

### Depletion of *COX11* alters the composition and activity of the respiratory apparatus in *P. patens*

In *P. patens*, COX11 is encoded by a single nuclear gene and the encoded protein sequence includes a putative mitochondrial targeting peptide (Supplementary Figure 2). Using publicly available transcriptomics data, we found that the *COX11* gene is expressed at significant levels in all tissues, although the highest transcript levels are detected in imbibed spores compared to other developing stages (Supplementary Figure 3). Three cysteine residues known to be involved in Cu^2+^ binding and essential for COX11 activity as a copper chaperone in yeast (Carr et al., 2002) are also conserved in Pp-COX11 (Supplementary Figure 2). All these observations are consistent with COX11 activity being conserved in *P. patens*.

**Figure 1.**
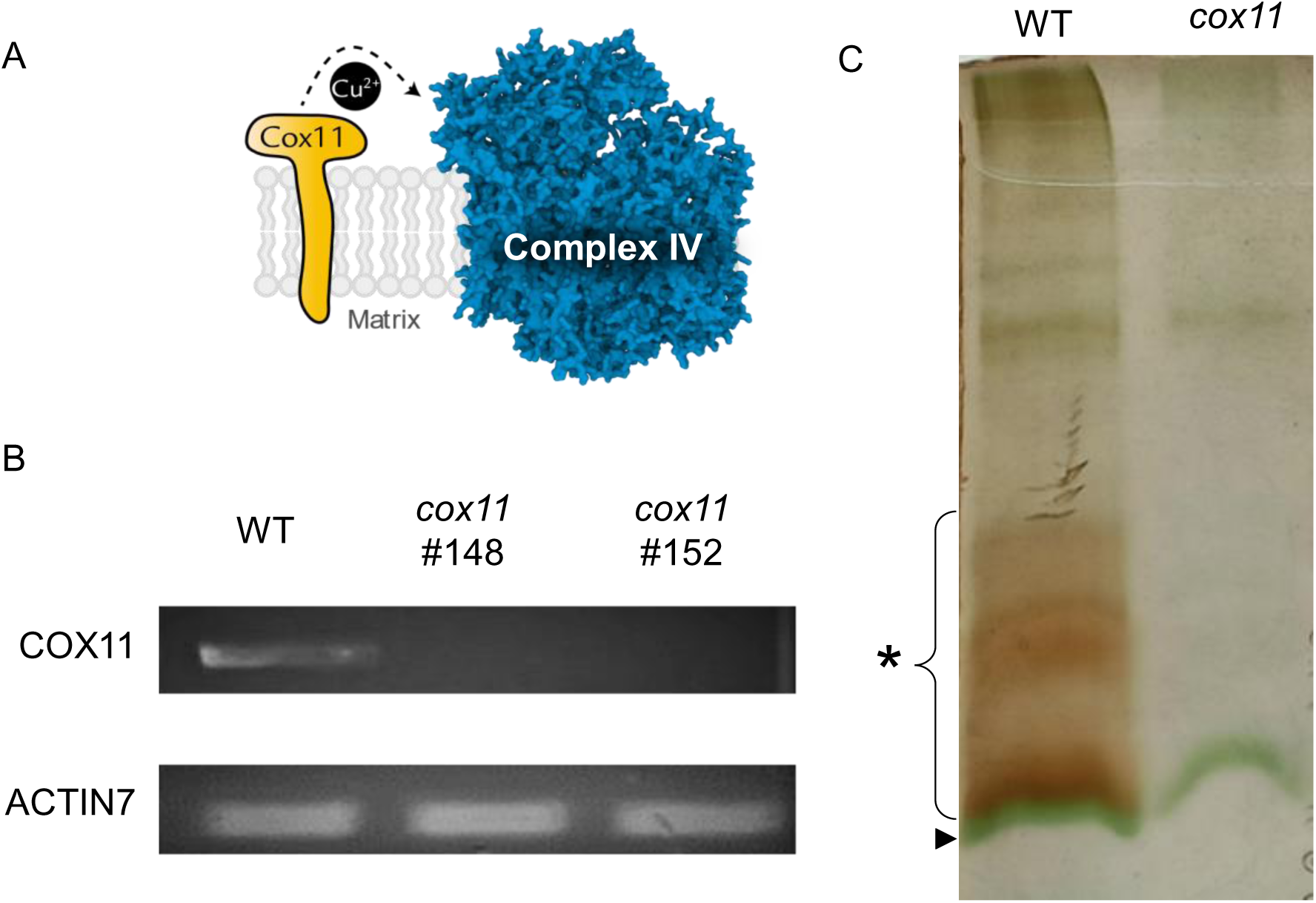
Isolation of knockout mutants for the gene *COX11* in *P. patens*. (A) Diagram showing the topology and function of COX11, which delivers Cu2+ into the active site of complex IV, essential for activity. The model of complex IV was retrieved from Maldonado et al., 2021, PDB entry 7JRO. (B) RT-PCR verification of the expression of *COX11* in the two independent lines #148 and #152. (C) In-gel activity staining after separation of crude membrane extracts by BN-PAGE. The arrow marks a band that corresponds to LHCII trimers (Järvi et al., 2011), and the asterisk marks the area with complex IV activity in the WT.

*P. patens cox11* mutants were generated by polyethylene glycol-mediated transformation, as schematized in Supplementary Figure 4. Stably hygromycin-resistant lines were validated by PCR to confirm the insertion in the expected genomic region (Supplementary Figure 4B). The absence of the *COX11* mRNA was confirmed by RT-PCR (Figure 1B). To verify that the loss of *COX11* led to a deficiency on CIV activity, crude membrane extracts, enriched in mitochondrial proteins, were separated by blue native polyacrylamide gel electrophoresis (BN-PAGE). After separation, in-gel activity assay for the cytochrome c oxidase showed no detectable activity in *cox11* plants, which was instead clearly detected in WT plants (Figure 1C).

### Growth is altered in *cox11* plants

All validated *cox11* lines showed severely impaired growth and delayed developmental phases (Figure 2). *cox11* plants remained longer than WT in the protonema phase and first gametophores were observed after 21 days, whereas in WT they were visible after 10 days (Figure 2; Supplementary Figure 5). Spore production could not be induced in *cox11* under the conditions normally effective for WT plants.

The growth phenotype of *cox11* was assessed under different conditions (Figure 2B). Addition of glucose and ammonia to the media had a positive effect on *cox11* growth, as it did for WT plants, and the respective difference remained (Figure 2B). Growth under continuous illumination caused a small but significant improvement in *cox11* growth compared to the control long day photoperiod. However, this increase was far from sufficient to rescue the growth difference with WT, which remained very large. Finally, growth under an elevated CO_2_ atmosphere, which induces carbon fixation and removes the negative effects of photorespiration, had only a minor positive effect on *cox11*, while strongly stimulating the growth of WT plants.

**Figure 2.**
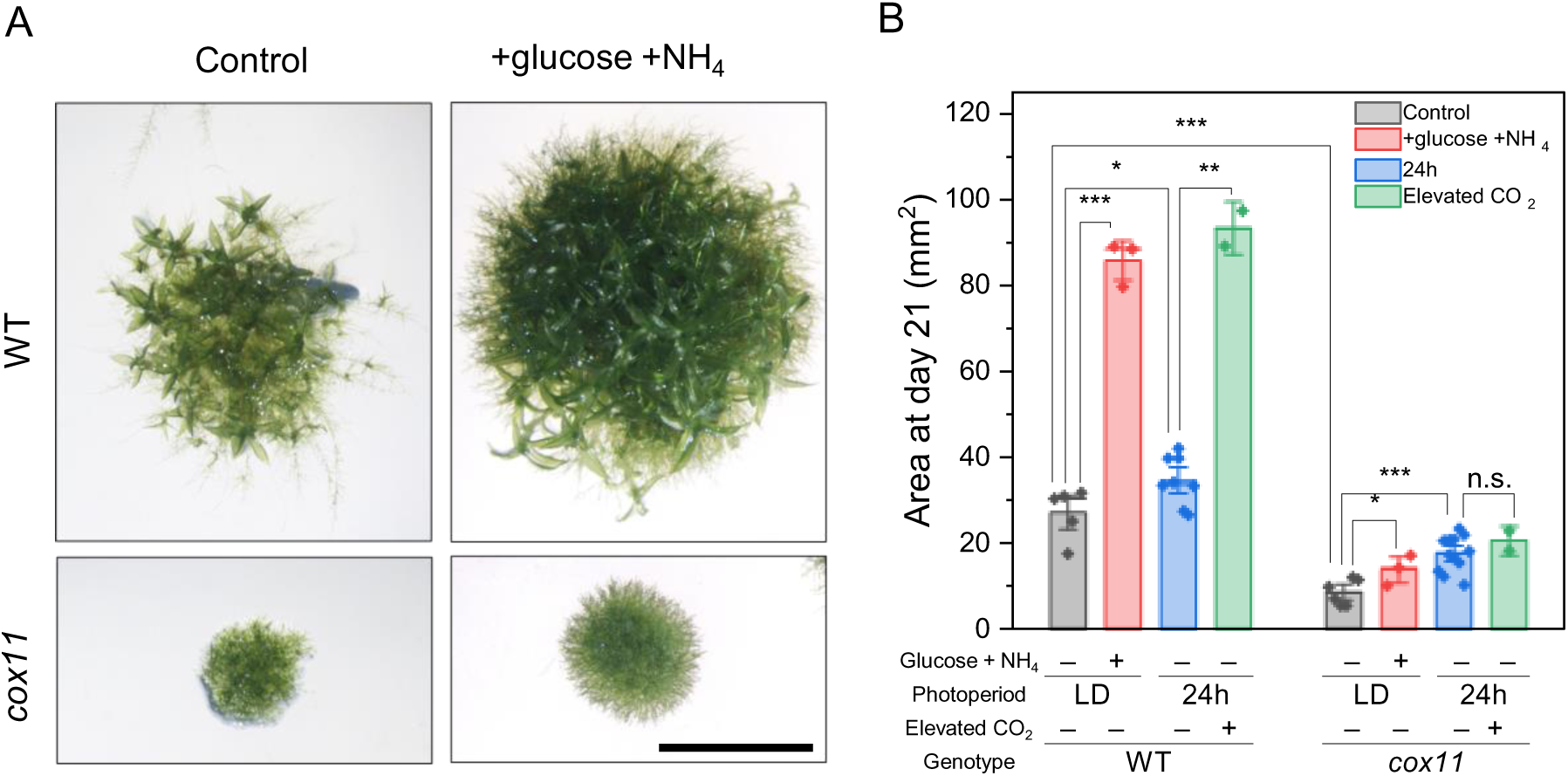
Growth phenotype of *cox11* mutants under different growth conditions. (A) Image of a representative colony of WT and *cox11* after 21 days of growth on solid PpNH4 medium. Scale bar is 5 mm. (B) Area quantification at day 21 of colonies grown under control conditions. LD, long day; 24 h, continuous illumination. Statistics: two-sample t-test, (***) p<0.001; (**) p<0.01; (*) p>0.05; (n. s.) not significant. *cox11* reports merged data from 2 independent lines.

### Composition and activity of the respiratory apparatus are altered in *cox11*

The effect of *COX11* depletion on the composition of the respiratory apparatus was verified by immunoblotting, using antibodies against some of the core proteins of the respiratory complexes (Figure 3A). The immunoblot analysis showed increased levels of the CI core subunit NAD9, the CII core subunit SDH2, and the ß-subunit of CV. Interestingly, the levels of the CIII MPP subunit were lower in *cox11* than WT plants. Protein levels of the alternative oxidase (AOX), which allows electrons to bypass the cytochrome pathway, were also lower in *cox11* than WT plants.

**Figure 3.**
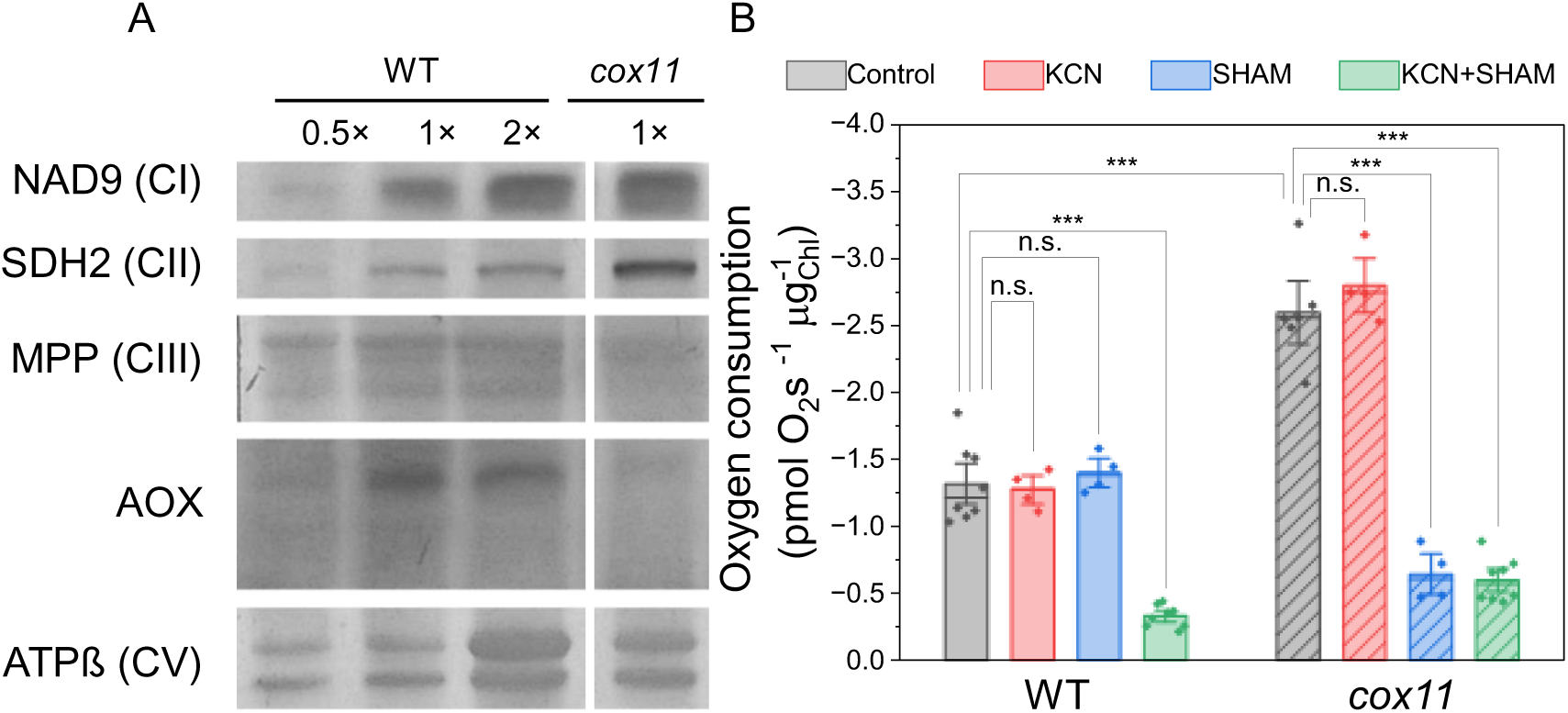
Alterations in the respiratory machinery of *cox11*. (A) Immunoblot of subunits of the different respiratory complexes. For complexes I, II, III and the AOX, total protein extracts were used. For complex V, crude mitochondrial extracts were used. Different amounts of proteins were loaded, expressed in multiples of the WT (0.5×, 1×, 2×). 1× corresponds to 2 μg of chlorophylls for NAD9, SDH2 and the AOX, 4 μg of chlorophylls for MPP and 30 μg of proteins for the ß-subunit. (B) Experiments of respirometry on intact protonema. Statistics: two-sample t-test, (***) p<0.001; (n.s.) p>0.05.

Respiratory activity was assessed by measuring O_2_ consumption in the dark in intact protonema tissues (Figure 3B). Interestingly, O_2_ consumption in the dark was enhanced in *cox11* lines, reaching approximately twice the activity of WT plants. This was independent of normalization to the amount of protein or chlorophylls, since the protein/chlorophyll ratio was unchanged between WT and *cox11*, with values of 90.7±4.0 (WT) and 90.8 ±0.6 (*cox11*) mg_prot_/mg_Chl_.

To better understand the cause of the increased O_2_ consumption in *cox11* plants, we repeated the measurements with the addition of chemicals that inhibit either the cyanide-sensitive or the cyanide-insensitive pathways, i.e., potassium cyanide (KCN) or salicylhydroxamic acid (SHAM), respectively. In WT, the effects of either KCN or SHAM on dark respiration were not significant when applied alone (Figure 3B), whereas when applied together they abolished virtually all the dark respiration activity (Figure 3B). This suggests that, when the cytochrome pathway is chemically blocked, the excess of electrons can be readily redirected through the AOX, which thus must have a very high capacity in *P. patens* WT. To confirm that this effect is not due to chemical specificity, we repeated the experiment using two alternative chemicals, antimycin A and n-propylgallate, and reached the same conclusions (Supplementary Figure 6A). In *cox11*, as in the WT, KCN alone had no effect on O_2_ consumption (Figure 3B). However, the addition of SHAM alone was able to completely inhibit O_2_ consumption (Figure 3B). This strongly suggests that the higher respiration in *cox11* is entirely due to an increased activity of the cyanide-insensitive, AOX-mediated pathway. The same effect was observed when the alternative AOX inhibitor n-propylgallate was used (Supplementary Figure 6B).

These results confirm that *cox11* plants lack a functional CIV, and that this loss completely compromises the cyanide-sensitive electron transport pathway. At the same time, the capacity of the cyanide-insensitive pathway is increased in *cox11*, and it is solely responsible for O_2_ consumption during dark respiration.

### Carbon fixation but not light conversion is altered in *cox11*

There is evidence that mitochondria are active in photosynthetic tissues during the day, and that they act synergically with chloroplasts to ensure an optimal photosynthetic performance (Hurry et al., 2005; Raghavendra et al., 1994; Shameer et al., 2019; Tcherkez et al., 2017; Vanlerberghe et al., 2020). Therefore, we evaluated whether the alterations in respiratory efficiency in *cox11* would have any effects on their photosynthetic performance. We quantified the abundance of different components of the photosynthetic machinery by immunoblotting and found no major differences (Supplementary Figure 7). We then quantified the photosynthetic activity of protonema pieces by measuring the rate of O_2_ evolution under saturating illumination, which induces maximal photosynthesis. In *cox11* the net photosynthetic activity was reduced (Supplementary Figure 8B) but this was attributable to the increased respiration, as gross photosynthesis was not different from the WT (Figure 4A), suggesting a good performance of the light reactions of photosynthesis.

Photosynthetic activity was further assessed by chlorophyll fluorometry (Figure 4B, C, D). The efficiencies of both photosystems I and II upon exposure to sub-saturating light, quantified from Y_I_ and Y_II_ respectively, were indistinguishable between WT and *cox11* (Figure 4B, C). The induction of non-photochemical quenching (NPQ), a photoprotective mechanism activated by a decreasing pH in the thylakoid lumen, was also unaltered (Figure 4D). Finally, the photosynthetic electron transport rate (ETR), quantified at steady state, was also indistinguishable between WT and *cox11* plants (Figure 4E).

The efficiency of the metabolic phase of photosynthesis was assessed by quantifying the CO_2_ fixation rate of protonema under a control light of 50 µmol photons m^-2^ s^-1^ (Figure 4F), which was lower in *cox11* compared to WT. These data indicate that the light phase of photosynthesis was not significantly altered in *cox11* mutants, unlike what was observed in the complex I deficient mutant *ndufa5* (Mellon et al., 2021), although net carbon fixation was slightly less efficient under moderate illumination.

**Figure 4.**
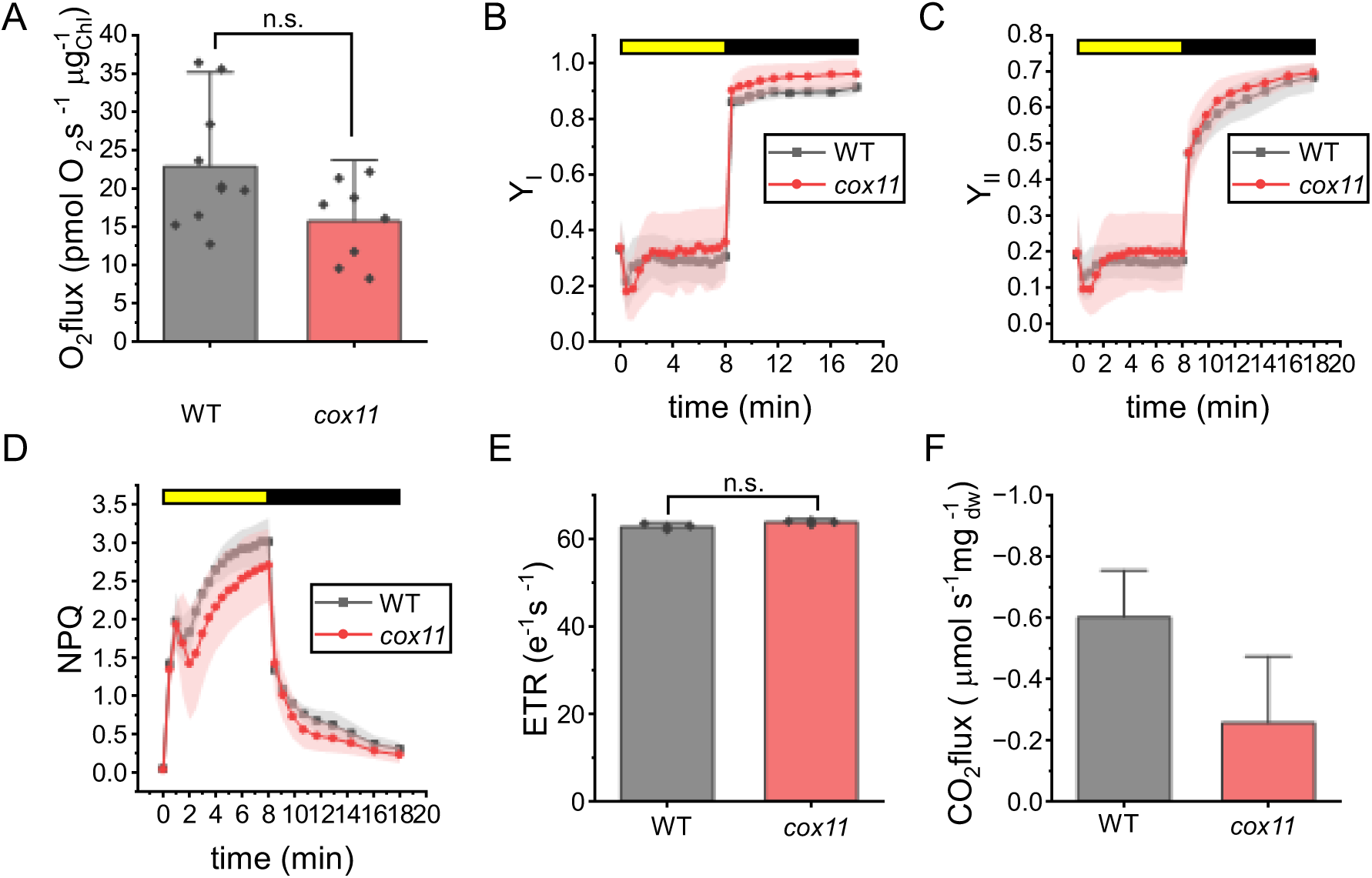
Evaluation of photosynthetic properties in *cox11*. (A) Gross evolution of O2 under saturating illumination. (B-D) The yield of PSI (YI, B), PSII (YII, C) and non-photochemical quenching (NPQ, D) were measured with Dual PAM 100 in plants exposed to 330 μmol photons m-2 s-1 of actinic light intensity. Yellow / black bars indicate when actinic light was on / off respectively. WT and *cox11* are shown respectively with black squares and red circles. Data are shown as average ± SD (n > 4). No statistically significant differences from WT plants were identified. (E) Electron transport rate of dark-acclimated plants grown under dim light, calculated from the ECS (electrochromic shift signal) after exposition to saturating light (300 µmol photons m-2 s-1) for 3-5 minutes. Activity was normalized to the total photosystem (PSI+PSII) content. Standard deviation is also reported (n > 6). (F) Net CO2 fixation rate under a control light of 50 µmol photons m-2 s-1. Error bars in A, E and F represent 1.5 times the SD. Statistics: two-sample t-test, (n.s.) p>0.05.

### The *cox11* transcriptome shows alterations in metabolism that are dependent from the photoperiod

In order to globally assess the impact of the mutation on metabolism, plant material from WT and *cox11* was used for transcriptome analysis by RNA sequencing. Since it could be expected that the impact of an altered mitochondrial activity could change during the day, depending on whether photosynthesis is active or not, plant material was harvested at different zeitgeber times (ZT), namely at the end of the night (ZT0), at the beginning (ZT2), or at the middle (ZT6) of the day (Supplementary Figure 9).

After comparing the lists of differentially expressed genes (DEGs) at each of the three timepoints tested, we classified the DEGs according to the time of the day they were altered, defining twelve different expression patterns (Figure 5, Supplementary Dataset 1). In all cases, the number of genes upregulated in *cox11* was larger than the number of downregulated genes. We performed pathway enrichment analysis on the DEGs at each of the three time conditions (Supplementary Figure 9C) and compared the pathways that were exclusive or shared between conditions (Supplementary Figure 10; Supplementary Dataset 2).

**Figure 5.**
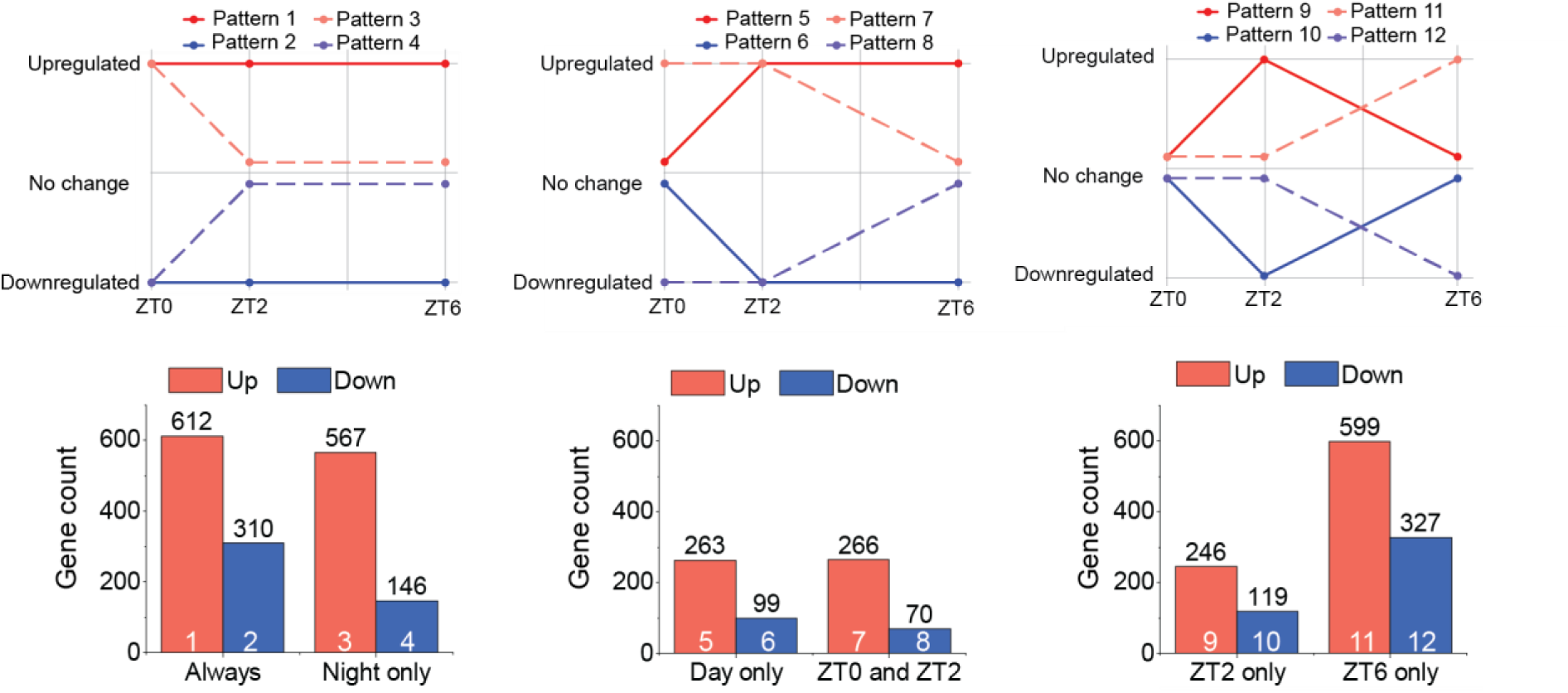
Overview of the diel regulation of the differentially expressed genes (DEGs) identified in *cox11*. We defined twelve different patterns and manually classified the genes accordingly (top). The number of genes included in each group is shown by the column charts (bottom). Numbers on top of bars show the number of included genes. Numbers in the bars identify the corresponding pattern. The lists of genes following each of the patterns are supplied as Supplementary Dataset 1C.

There were 612 genes significantly upregulated genes in all conditions (Pattern 1), although we did not identify any specific pathway that were always significantly upregulated. On the other hand, we identified 310 genes and 46 pathways that were always downregulated in *cox11* as compared to WT (Pattern 2; Supplementary Figure 10A). Several of these genes are involved in cell wall biogenesis and remodelling, or metabolism of structural polysaccharides, suggesting alterations in cell wall architecture (Le Gall et al., 2015). The expression of various transporters was also reduced in *cox11*, including transporters of amino acids, inositol, ammonium, phosphate, or zinc, as well as several aquaporins. This could reflect a reprogramming of the metabolic machinery of the cell to cope with a strongly reduced growth, which necessarily reduces cell wall synthesis and elongation as well as nutrient demand (Boursiac et al., 2008; X. Liu & Bush, 2006; Vandeleur et al., 2014; Yang et al., 2020).

713 genes showed differential expression (146 down, 567 up) only during the night, but unchanged levels during the day. The upregulated pathways at night (Pattern 3, Supplementary Figure 10B) included a group of transaminases involved in the metabolism of amino acids, suggesting that nitrogen metabolism might be altered in *cox11*, particularly at night. Following Pattern 3 we also detected several genes associated with stress responses that included many different types of heat shock proteins (Hsp), either from the small (Hsp17.4, Hsp17.6, Hsp20) or large (Hsp90) families (Waters & Vierling, 2020), as well as proteins of the BAG family, which are likely chaperone regulators and might assist heat shock proteins in their chaperone role (Doong et al., 2002; Kabbage & Dickman, 2008).

The induction of heat shock proteins could be part of a more general response to the accumulation of unfolded proteins, which would trigger the unfolded protein response in the endoplasmic reticulum (UPR^ER^). However, the expression levels of some genes previously defined as markers of UPR^ER^ in *P. patens* (Lloyd et al., 2018) were not significantly higher in *cox11* (Supplementary Table 4), therefore not supporting a general induction of UPR^ER^ in *cox11*. We also checked the expression levels of genes encoding antioxidant enzymes, but none was significantly upregulated in *cox11* plants (Supplementary Table 5), suggesting that plants were not suffering of general oxidative stress. The only exception was a superoxide dismutase, PpFSD2, predicted to be targeted to the chloroplast (Higashi et al., 2013, Supplementary Table 5), which was always induced in *cox11* plants. Therefore, the expression pattern of the different heat shock proteins suggests that a stress response is induced during the night, but it is then alleviated during day, likely by exposure to light, which apparently can relieve the stress. The induction of heat shock proteins is likely not due to oxidative stress, because antioxidants did not get induced.

The genes repressed by night (Pattern 4; Supplementary Figure 10B) included one of the three annotated genes encoding nitrate reductase isoforms in *P. patens*, a key enzyme for nitrogen assimilation (Chamizo-Ampudia et al., 2017; Medina-Andrés & Lira-Ruan, 2012) whose activity can be regulated during abiotic stress (Fu et al., 2018). In addition, there was a group of 35 transcription factors (TFs) of different families, including apetala 2 (AP2), MYB, MYC, ABA-inducible, and GRAS. Some of these transcription factors retained their regulation also during the first hours of day (Pattern 8, Supplementary Figure 10D), but all of them recovered normal values at ZT6. AP2, with 149 members in *P. patens*, has been linked to defence and stress responses, growth and development (Aoyama et al., 2012; Ishikawa et al., 2019; Z. S. Xu et al., 2011; J. Zhang et al., 2020). In turn, MYB factors have been linked to protonemal growth and the transition from chloronema to caulonema (Pu et al., 2020), and GRAS factors to the development of gametophores and spores (Beheshti et al., 2021). Overall, the altered levels of these TFs are most likely correlated with the delay in development program observed in *cox11*.

We found gene sets whose expression was normal at night but changed during the day. Most of those upregulated only at the beginning of the day (Pattern 9 Supplementary Figure 10E) encoded chloroplast proteins, including starch synthases, required for carbon storage (Pfister & Zeeman, 2016); arogenate dehydratases, required for biosynthesis of the amino acid phenylalanine (C. Bonner & Jensen, 1987; Maeda & Dudareva, 2012); and ATP/ADP transporters, which might be linked to the inter-compartment trafficking of energy equivalents. On the other hand, the transcripts found at higher levels throughout the day (Pattern 5; Supplementary Figure 10C) included 6 genes related to starch biosynthesis, either starch synthases or ADP-glucose synthases, which catalyse the committing step of starch biosynthesis (Pfister & Zeeman, 2016; Stitt & Zeeman, 2012), and several genes involved in ribosome biogenesis, transcription, and translation suggesting that the gene expression and protein synthesis rates are increased in *cox11*, particularly during day.

Some genes were instead repressed only at the beginning of the day (Pattern 10, Supplementary Figure 10E), including two putative urea transporters (Pp3c14_7160 and Pp3c1_21890), and enzymes required for amino acid or glucose metabolism, including many enzymes of glycolysis. This observation further strengthens the previously proposed hypothesis that *cox11* plants present altered central metabolism at night and at the night-to-day transition, affecting both N and C metabolism.

### Metabolomics and integrated pathway analysis show extensive alterations in carbon and nitrogen metabolism in *cox11*

To further investigate the metabolic alterations in *cox11* shown by the transcriptomics data, we performed untargeted metabolomics on the same samples used for transcriptome analyses. We uniquely identified a total of 65 compounds, 23 by GC-MS and 42 by IC-MS, including most of the primary amino acids except for arginine, cysteine, histidine, lysine, proline and tryptophan; several monosaccharides; intermediates of glycolysis and TCA cycle; CBB cycle intermediates; and nucleotides, among other compounds (Figure 6A; Supplementary Dataset 3). Pairwise comparisons of samples at each zeitgeber time identified 34 metabolites at significantly higher or lower levels in one or more conditions.

As we observed from the transcriptome, the largest number of differences in the *cox11* metabolome emerged when comparing the end of night and early morning samples. Overall, the metabolomic data are consistent with the transcriptome in showing important changes in carbon metabolism (Figure 6B). Seven out of the ten intermediates of the glycolytic pathway were identified as altered in our experiments and, except for glucose-6-phosphate and fructose-6-phosphate, all of them accumulated at lower levels in *cox11* at ZT0. The transcript levels of the pathway enzymes did not show a consistent regulation, as most of them were unchanged while a few isoforms were either upregulated or downregulated. Similarly, four of the eight intermediates of the TCA cycle were detected by our metabolomics approach (citrate, 2-ketoglutarate, fumarate and malate), and all except fumarate were lower. Transcript levels of the TCA enzymes were mostly unaltered, except for the repression of an annotated isoform of malate dehydrogenase (Pp3c4_20940) and a slight induction of some citrate synthase and succinate dehydrogenase isoforms. Globally, metabolomic data strongly suggest that carbon catabolism was blocked or slowed down in *cox11* at night (Figure 6B). Interestingly, this alteration was no longer observed during the day, when intermediates returned to their normal values despite a more general transcriptomic activation (Figure 6C, D). Notably, at ZT2 we observed the accumulation of both ribulose-5-phosphate and ribulose-1,5-bisphosphate, the carbon skeleton required for RuBisCO to fix CO_2_ and its direct precursor; this is consistent with the reduced rate of carbon fixation that we reported previously (Figure 4F).

**Figure 6.**
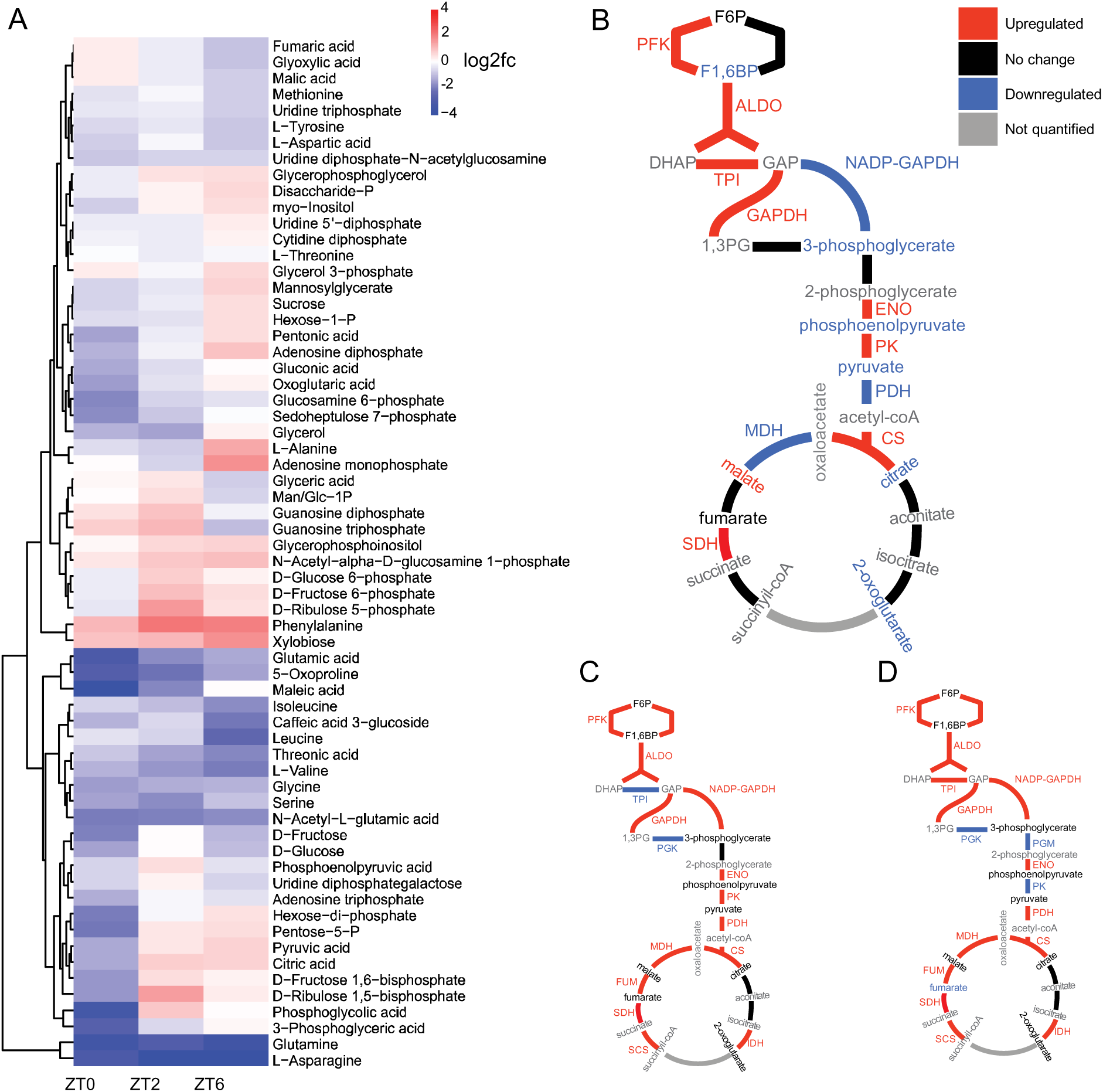
Overview of metabolomics data. (A) Heatmap showing hierarchical clustering of compounds. (B-D) Representation of metabolic pathway of glycolysis, pyruvate dehydrogenase and tricarboxylic acid cycle integrating transcriptomics and metabolomics data for *cox11* at ZT0 (B), ZT2 (C) and ZT6 (D). Enzyme names: PFK, phosphofructokinase; ALDO, aldolase; TPI, triose phosphate isomerase; NADP-GAPDH, NADP-specific glyceraldehyde dehydrogenase; GAPDH, NADH-dependent glyceraldehyde dehydrogenase; PGK, phosphoglycerate kinase; PGM, phosphoglycerate mutase; ENO, enolase; PK, pyruvate kinase; PDH, pyruvate dehydrogenase; CS, citrate synthase; IDH, isocitrate dehydrogenase; SCS, succinyl-CoA synthetase; SDH, succinate dehydrogenase; FUM, fumarase; MDH, malate dehydrogenase. (C) Metabolic pathway of glycolysis, pyruvate dehydrogenases and TCA cycle showing transcript and metabolite regulation in *cox11* at ZT0.

Not only carbon metabolism, but also nitrogen metabolism was clearly altered in *cox11* plants. Phenylalanine was the only proteinogenic amino acid identified at higher levels in *cox11* at the beginning of the day, whereas the levels of the other detected amino acids were reduced in *cox11* at ZT0 only (glycine), at ZT0 and ZT2 (asparagine, glutamate, serine, valine) or always (glutamine) (Figure 6A, Supplementary Dataset 3). The lower levels of glutamate, glutamine and asparagine are particularly significant since they are among the main metabolites involved in nitrogen assimilation from inorganic nitrogen sources in plants (Gaufichon et al., 2010; Liao et al., 2022; Singh, 1998). The glutamate/glutamine and aspartate/asparagine ratios were also altered (Supplementary Figure 11), further suggesting a reduction in nitrogen assimilation efficiency. Arginine, with its high nitrogen-to-carbon ratio, is also used for nitrogen storage and mobilization in plants (Winter et al., 2015; Yoneyama & Suzuki, 2020). We could not detect arginine in our metabolomic analysis, but we found the arginine biosynthetic machinery to be transcriptionally repressed. Furthermore, N-acetyl-L-glutamate, a substrate for glutamine-regulated arginine biosynthesis (Chellamuthu et al., 2014; Slocum, 2005), was always found at lower levels in *cox11*.

All these observations are consistent with *cox11* cells modulating their metabolism towards a starvation situation by blocking the energetically costly process of nitrogen assimilation at night. Nitrogen assimilation and mobilization are therefore altered in *cox11*, especially at night, and potentially affecting a plethora of metabolic and cellular processes.

### Starch mobilization is impaired and energy availability is unchanged in *cox11*

By observing micrographs of moss samples by transmission electron microscopy (TEM), we found that *cox11* accumulates large starch granules in their chloroplasts. Even after 16 hours of incubation in the dark, while starch granules were absent or relatively small in WT plants, they were large and occupied most of the chloroplast area in *cox11* (Figure 7A). The quantification of starch content at ZT6 indeed confirmed that *cox11* had more than twice the amounts of starch compared to the WT (Figure 7B). To determine whether the starch degradation capacity of *cox11* was altered, we separated total *cox11* extracts under native conditions in gels containing solubilised potato starch and estimated total and maximal amylolytic activity by in-gel activity. This test clearly shows that amylolytic activity was higher in *cox11* than in WT plants (Figure 7C). In addition, at least one more band appeared in *cox11*. These data suggest that the hydrolytic activity in *cox11* was not reduced but actually increased compared to WT. Therefore, starch accumulation cannot be explained by a reduced capacity of amylolytic activity and most likely involves an increased rate of synthesis and/or reduced degradation *in vivo* due to the blockade of downstream metabolic processes such as glycolysis.

**Figure 7.**
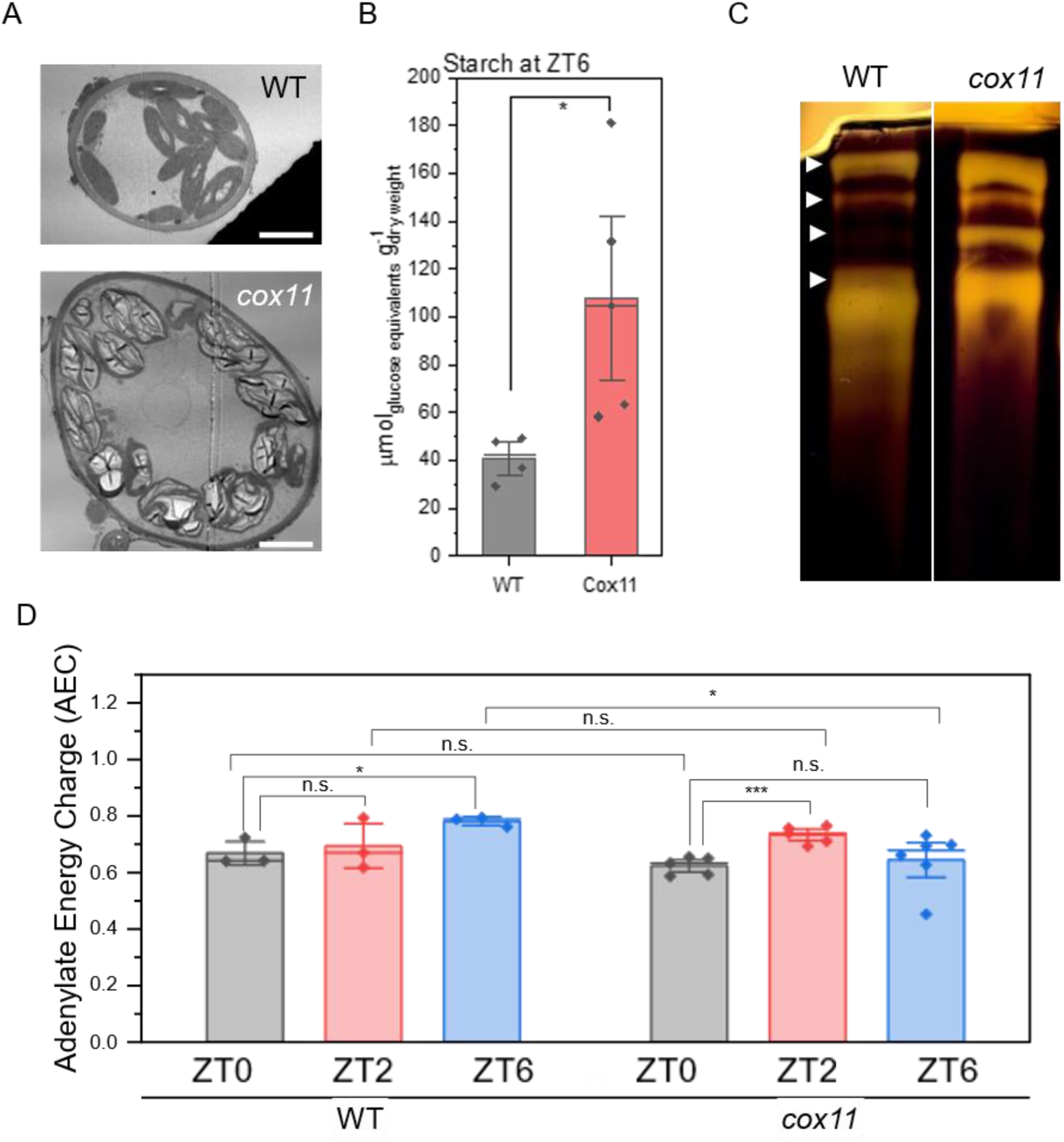
Starch accumulation and energy availability in *cox11*. (A) Representative micrographs of WT and *cox11* cells showing differences in the amount and dimension of starch granules inside of chloroplasts. Plant samples were fixed after 16 hours of darkness. Scale bar is 5 µm. (B) Quantification of starch in total extracts harvested at ZT6. (C) In-gel activity of starch degrading enzymes. White arrow heads show the four main identified bands of activity. It is not possible to identify the enzymes responsible for each amylolytic activity due to the lack of literature data on amylases from *P. patens*. (D) Relative adenylate energy charge (AEC) of WT and *cox11*. Statistics: two-sample t test. (***), p<0.001; (*), p<0.05; (n.s.), p>0.05.

The previously reported deficiency of glycolysis intermediates, together with the accumulation of starch, suggest a deficient mobilization of carbon stores during the night in *cox11*. A way to measure the amount of energy available for the cell is to quantify the adenylate energy charge (AEC), based on the levels of adenosine-5’-tri-, -di- and -monophosphates (ATP, ADP, AMP) (Atkinson, 1968). We used adenylate levels from untargeted metabolomics to calculate the relative AEC values for both WT and *cox11* plants (Figure 7D; see Material and methods section for calculation details). We found that AEC values in *P. patens* WT plants ranged between 0.6 and 0.8, a range consistent with previous experiments in other plants (Hampp et al., 1982; Lange et al., 2008). In WT, the AEC increased throughout the day, with the difference becoming significant at ZT6. In *cox11*, the value at ZT0 was not significantly different from that of the WT; then, we observed an increase at ZT2 similar to that of WT, but thereafter the AEC dropped almost to ZT0 values as the day progressed (Figure 7D). Even if some alterations are indeed present, this data suggests that *cox11* plants do not suffer from major energy starvation.

### External supply of amino acids partially rescues the growth phenotype of cox11 plants

Given the general alteration in amino acids levels and metabolism, we investigated whether the external application of amino acids could rescue the growth phenotype of *cox11* by providing metabolic intermediates that could restore one or more essential metabolic pathways that were impaired in *cox11* under control conditions. Therefore, we compared the growth of *cox11* plants grown under control conditions, i.e., where the sole nitrogen source was inorganic nitrate, or in enriched media where the nitrogen source was both inorganic nitrate and organic nitrogen in the form of one type of amino acid.

On WT plants, all the amino acids except for glutamate had a positive or negative significant effect on growth that could be noticed during the first 2-4 weeks of culture in most cases (Supplementary Figure 12A). All the amino acids that reduced the growth rate of the WT also had a negative effect on *cox11*, except for phenylalanine which did not cause a growth reduction on *cox11*. For some of these (histidine, isoleucine, lysine, proline), the negative effects on *cox11* were no longer observed after seven or more weeks of culture.

The amino acids that had a positive effect on WT (alanine, arginine, asparagine, aspartic acid, glutamine, glycine and serine) were also beneficial for *cox11*, and in some cases the growth improvement of *cox11* was enough to rescue the growth impairment associated with the mutation (Supplementary Figure 12B). Serine is reported as example of amino acid with a strong beneficial effect (Figure 8) and its addition not only increased the growth area of the plant, but it also induced an earlier formation of phyllids and gametophore-like structures on *cox11* (Figure 8B), suggesting a rescue of the developmental program as well. The same was also observed for glycine, which is not surprising considering that serine and glycine are readily interconvertible (Supplementary Figure 12B).

We demonstrate that the external addition of several amino acids can partially rescue the phenotype of *cox11*, which is consistent with *cox11* having a globally altered metabolism in which some metabolites are lowly available, compromising the function of anaplerotic reactions required for the proper use of energy and resources.

**Figure 8.**
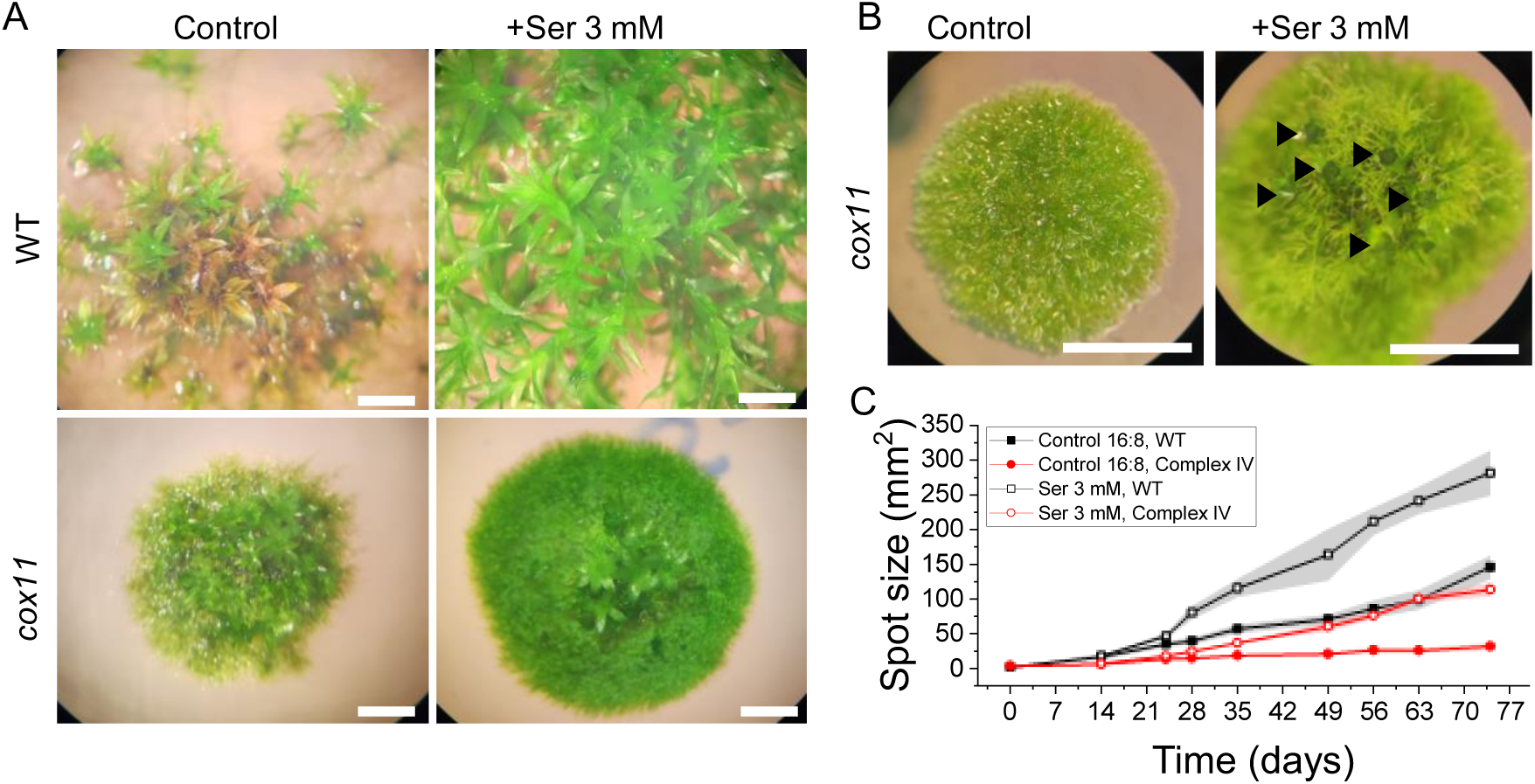
Effect of external supply of serine on growth. (A) Images of moss colonies after 42 days of growth on control medium or medium containing 3 mM serine. Scale bars are 2 mm. (B) Detailed view of 21 days old *cox11* plants grown in media with or without the addition of 3 mM serine. Black arrows mark developing gametophores. Scale bars are 2 mm. (C) Growth curve of *cox11* supplemented with 3 mM serine.

## Discussion

### *cox11* plants lack a functional cytochrome c oxidase

Here we report the isolation and characterization of mutant lines of the moss *Physcomitrium patens* knocked out for the conserved copper chaperone COX11. COX11 has been reported to play an essential role in the biogenesis of the mitochondrial complex IV (cytochrome c oxidase), where it is required for the insertion of copper ions into the active site of COX2 (Meyer et al., 2019). Its depletion leads to a complete inactivation of complex IV in yeast (Banting & Glerum, 2006; Carr et al., 2002; Tzagoloff et al., 1990). Accordingly, *cox11* knockdown lines in Arabidopsis showed a strong reduction in complex IV activity (Radin et al., 2015), but the complete knockout was not viable (Radin et al., 2015).

In this work, we characterize plants completely depleted of COX11, which confirm that this protein is essential for CIV activity also in the moss *P. patens*, as crude membrane extracts of *cox11* plants had undetectable levels of cytochrome c oxidase activity. To our knowledge, only one other plant mutant has been reported to have undetectable levels of cytochrome c oxidase, the *cod1* mutant in Arabidopsis (Dahan et al., 2014). However, *cod1* plants suffered from general and severe alterations and could only be analysed by rescuing immature seeds, and no analysis under physiological conditions was possible (Dahan et al., 2014).

The isolation of a viable *cox11* mutant was possible in *P. patens* because this plant is photoautotrophic throughout its development, allowing to avoid heterotrophic tissues, such as roots (Sakakibara et al., 2003). Mutants can be isolated by vegetative propagation under continuous light, by-passing sexual reproduction and spore formation/generation, thus avoiding life cycle stages where photosynthesis is not active and thus respiration essential. In addition, *P. patens* cells are haploid during most of their life cycle, allowing the phenotype associated with any genetic modification to be directly assessed without reaching homozygosity. In *P. patens* the effects of inactivation of mitochondrial respiration can thus be assessed in fully photosynthetically active cells, providing a highly valuable model for studying the physiological consequences of the constitutive blockade of the cyanide-sensitive respiratory pathway in plants.

### *cox11* has altered mitochondrial respiratory chain composition and activity

By monitoring the rate of oxygen consumption, we demonstrated that the cyanide-sensitive pathway is completely inactivated in *cox11* plants, which entirely rely on the AOX-mediated alternative pathway to maintain the ubiquinone pool in the oxidized form. Such a prominent activity of the alternative pathway implies that the only complex available for proton translocation to fuel oxidative phosphorylation in *cox11* mitochondria is complex I. Interestingly, the oxygen consumption rate is increased in *cox11*, demonstrating that the AOX pathway has a high electron flux capacity. This is observed even in *P. patens* WT plants where AOX can sustain strong respiratory activity in the presence of cyanide. *cox11* mutants also show increased protein levels of NAD9, a core subunit of complex I, and SDH1, the catalytic subunit of complex II, suggesting an increased capacity for electrons to enter the modified respiratory chain, that, together with AOX, enabled the observed increased O_2_ consumption.

AOX protein levels were not higher in the mutant, but this protein is known to be highly post-translationally regulated (Ho et al., 2008; Moellering & Benning, 2010; Rhoads et al., 1998) and thus increased activity could be achieved without stronger protein accumulation. Consistent with this observation, Arabidopsis plants with reduced levels of cytochrome c and complex IV also showed unaltered or even reduced levels of Aox1a, the homolog of Pp-Aox (Florez-Sarasa et al., 2021). Remarkably, plants with altered complex I activity, including *P. patens*, commonly show AOX overaccumulation, which is not the case here (Fromm, Senkler, et al., 2016; Gutierres et al., 1997; Mellon et al., 2021; Meyer et al., 2009; Pétriacq et al., 2017; Sabar et al., 2000). AOX overaccumulation in these plants, however, is associated with the induction of mitochondrial retrograde signalling (Giraud et al., 2009; Millar et al., 2011), which is likely not occurring in *cox11* plants here.

The transfer of electrons from complex I to molecular oxygen through the alternative oxidase is much less efficient than the cytochrome pathway in ATP production, but it nevertheless ensures the consumption of reducing power, which is likely a pivotal function in photosynthetically active cells (Dahal et al., 2017; Krämer & Kunz, 2021; Moreno-García et al., 2022). It is well known that the photosynthetic electron transport produces NADPH in excess of ATP for CO_2_ fixation, and multiple mechanisms exist to balance this ratio, such as cyclic and pseudo cyclic electron transport (Shikanai & Yamamoto, 2017). Mitochondrial respiration is probably a major contributor in balancing the extra reducing power in the cell (Burlacot & Peltier, 2023), as it has been suggested from experiments with inhibitors (Yoshida et al., 2006) or modelling analyses (Shameer et al., 2019). In support of this idea, increasing the flow of reducing equivalents from chloroplasts to mitochondria by overexpressing the protein PAP2 had a positive effect in balancing the NADPH/ATP ratio in chloroplasts and increased the efficiency of photosynthesis in Arabidopsis (Voon et al., 2021).

It is interesting to observe that, unlike complex I mutants, *P. patens cox11* plants did not show a major alteration of photosynthetic electron transport reactions, suggesting that the altered respiratory apparatus is indeed able to sustain the consumption of reducing power sufficient to balance the photosynthetic activity, not inducing any secondary regulation, as shown by the substantial similarity of the activity of the light phase of photosynthesis.

### Carbon metabolism is drastically affected in *cox11*

*cox11* plants could be propagated vegetatively and were viable, but their growth was severely impaired. Remarkably, this is the case even if all cells of protonema, the tissue that we used for our experiments, are fully photosynthetically active and even when plants are grown under 24 hours of illumination, ensuring a constant supply of energy from photosynthesis and eliminating the potential effects of lack of energy during the night. This contrasts with respiratory mutants depleted in complex IV in the green alga *Chlamydomonas reinhardtii,* which showed a growth delay only when grown under hetero/mixotrophic conditions but were not significantly different from the WT in minimal medium when photosynthesis was active (Colin et al., 1995). This suggests that the role of respiration in photosynthetic organisms changed during plants evolution (Mellon et al., 2021).

We have shown that *cox11* plants have an increased electron transport rate that is entirely due to a more active cyanide-insensitive pathway. Such an increase could enable *cox11* plants to maintain a similar or only slightly lower levels of ATP via oxidative phosphorylation compared to the WT. This is supported by the observation that the relative adenylate energy charge (AEC) does not show significant different in *cox11* even at night. The reduced growth of *cox11*, and consequent reduction in energy consumption, might also contribute to keeping the AEC close to normal values. A similar effect has been shown in tobacco plants depleted in complex I activity, where the remodulation of electron transport increased the efficiency of oxidative phosphorylation, so that the predicted ATP production was maintained (Vidal et al., 2007). More recently, we similarly observed in *P. patens* complex I mutants increased chloroplast ATPase activity to maintain ATP leaves (Mellon et al., 2021).

*cox11* plants are substantially unaltered in the light reactions of photosynthesis and thus are able to exploit light energy for synthesis of ATP and NADPH. Despite a lower rates of carbon fixation, they also accumulated more starch, which is a product of photosynthesis confirming their ability to perform photosynthesis and suggesting that this pathway could not be the cause of the growth defects described in *cox11*. The elimination of photorespiration by growing plants under elevated CO_2_ also did not have any positive effect on growth. The cultivation under continuous illumination also did not show rescue of the phenotype, all observations that support the hypothesis that mutants are not in energy deficiency.

The synthesis of starch was also upregulated at the transcript level, by the induction of several starch synthases during the day, again consistent with the idea that photosynthesis and carbon fixation activity were functional and probably in excess with respect to metabolic consumption. The fact that amylolytic activity of *cox11* extracts was higher suggests that the observed starch over-accumulation in *cox11* is not due to an impairment of its mobilization but rather to a limited ability to consume the products of its degradation. Since starch degradation is activated during the night, when mitochondrial electron transport is the main pathway for energy generation, it is likely that its impairment will affect sugars metabolization and consequently starch degradation, despite amylolytic activities being even induced. Consistent with this conclusion, a higher starch accumulation has been reported for Arabidopsis mutants with decreased activity of the cyanide-sensitive pathway (Racca et al., 2018). These observations all consistently point to the fact that photosynthesis and carbon fixation are all active. Carbon is also stored in form of starch, but cells have impaired ability to mobilize the reduced carbon available, as evidenced by the reduced levels of glycolytic intermediates at night when the catabolism of starch should provide the cell with monosaccharides to fuel respiration.

Consistent with this hypothesis, attempts to rescue the growth phenotype by inducing a more favourable energetic state largely failed. Neither additional light of any duration or intensity nor increased CO_2_ supply were able to rescue the phenotype, suggesting that photosynthesis, CO_2_ fixation, or photorespiration, even if affected by the mutation, were not the main responsible for the growth impairment. The addition of glucose and ammonium tartrate had a positive effect on growth, but this was that same as in WT and the large growth defect of the mutant was maintained. In summary, the growth defects of *cox11* could not be attributed to a simple energy deficit, but rather to an insufficient capacity to utilize the reduced carbon synthesized thanks to photosynthesis.

### Mitochondrial metabolism has an essential impact on plants amino acid biosynthesis

The blockade of respiration had a major impact on amino acid metabolism, as evidenced by both transcriptomic and metabolomic data. Glycolysis and the TCA cycle are pivotal for the plants cell as they provide not only readily available energy in form of nucleotide phosphates such as ATP and GTP, but also metabolic intermediates for anaplerotic reactions (Sweetlove et al., 2010; Y. Zhang et al., 2018). For example, glutamate, found at lower levels at night in *cox11*, is an important bridge between carbon and nitrogen metabolism because it can be reversibly converted to the TCA intermediate 2-ketoglutarate (Forde & Lea, 2007; Hodges, 2002). The shortage of glycolysis and TCA cycle intermediates is therefore expected to alter the processes of nitrogen assimilation and mobilization. Indeed, nitrate assimilation in leaves of vascular plants, can occur at night by using stored carbohydrates (Yoneyama et al., 1987; Yoneyama & Suzuki, 2020).

Ammonium assimilation is done mainly through the glutamine synthetase/glutamate synthase (GS/GOGAT) cycle, although it could also possibly be assimilated via glutamate dehydrogenase (GDH) (Singh, 1998). The GS/GOGAT cycle requires glutamate, glutamine, and 2-ketoglutarate, all of which were found at lower levels in *cox11* at night (Gaufichon et al., 2010; Yoneyama & Suzuki, 2020). Nitrogen assimilation proceeds in part through the amination of aspartate and glutamate into asparagine and glutamine, respectively (Yoneyama et al., 1987). Not only the absolute values of the aminated forms asparagine and glutamine were lower, but also the aspartate/asparagine and glutamate/glutamine ratios were much higher in *cox11*, especially during the day, i.e. the deaminated forms of the amino acids were predominant, suggesting that nitrogen assimilation was not efficient.

These alterations of amino acids content and biosynthetic pathways is consistent with the growth phenotype of *cox11* plants on supplemented media. Even if there was no optimization of the concentrations employed, amino acid treatments generally had beneficial or at least neutral effects in *cox11*, even in cases where they had a toxic effect on WT growth, a phenomenon commonly reported in plants (C. A. Bonner et al., 1992, 1996; Forsum et al., 2008). In some cases, the amino acids external supply showed a particularly strong improvement of growth and significant rescue of the gap with WT. This is true for asparagine, glutamine, serine, three amino acids detected at lower levels at ZT0 and ZT2 in *cox11*. The most significant improvement was achieved when plants were grown in the presence of the amino acid serine, whose positive effect is not attributable to its involvement in photorespiration, since growth under high CO_2_ did not have a positive effect on *cox11* plants.

These data thus suggest that respiration plays a key role at the interface between carbon and nitrogen metabolism. When inactivated, cells are unable to support the effective incorporation of carbon skeletons into amino acids, which severely impairs their biosynthesis and thus has a major negative impact on growth.

## Material and methods

### Plant material and growth conditions

*Physcomitrium patens* (Gransden ecotype) was amplified through vegetative propagation as done previously on solid PpNH_4_ medium (Mellon et al., 2021). If not stated diversely, physiological and biochemical characterizations were performed on 10 day-old tissue cultivated in PpNO_3_ medium (Mellon et al., 2021) at 22 °C and a long-day photoperiod (light:dark 16h:8h).

### Generation of *COX11* knockout lines

For generation of *COX11* knockout (KO) lines, up- and downstream regions of the locus harbouring the *COX11* gene were cloned into a BHRf plasmid, which carries a hygromycin resistance cassette (Supplementary Figure 1A). The construct was linearized with the restriction enzyme PvuII (Thermo Fisher Scientific) and used for gene targeting through polyethyleneglycol (PEG)-mediated transformation as described previously (Mellon et al., 2021). Genomic DNA (gDNA) was extracted from stably resistant clones following a fast extraction protocol with some modifications (Edwards et al., 1991) and PCR amplifications of recombination cassette were performed on extracted gDNA. Protonemata of *P. patens* grown on PpNH_4_ were homogenized in 2 mL tubes using 3-mm zirconium glass beads (Merck) in presence of 500 µL of cold TEN buffer (Tris-HCl 100 mM, pH 8.0; EDTA 50 mM; NaCl 500 mM). After the addition of 35 μL of SDS 20 %, samples were incubated at 65 °C for 5 minutes. Then, 130 μL of potassium acetate 5 M were added, samples were kept on ice for 5 minutes and centrifuged at 4 °C for 10 minutes at 13,000 g. The supernatant was transferred to a clean 1.5 mL tube containing 500 μL of isopropanol at -20 °C, mixed by inversion and incubated at -20 °C for 10 minutes. Then, a series of centrifuges at 4 °C and 13,000 g were performed. Supernatant was discarded after a first centrifuge of 10 minutes, and the pellet was suspended in 500 μL of ethanol 70 %. The resultant pellet of a second centrifuge of 5 minutes was suspended in 150 μL of ethanol 70 %. A last centrifuge of 2 minutes was performed and the resultant pellet was dried off under the chemical fume hood and suspended in 50 μL of water. This DNA solution was kept at -20 °C until used.

To confirm that *cox11* lines lacked the *COX11* expression, reverse transcription (RT)-PCR was performed on cDNA using RevertAid reverse transcriptase (Thermo Fisher Scientific) synthesized after RNA extraction. Primers used for construct design and line validation are included as Supplementary Table 1.

Three independent lines, named #148, #150 and #152, were used in the experiments reported in this article. Each experiment contains data obtained from at least two independent lines.

### Public transcriptomic data and sequence alignment

The expression levels of *COX11* in different tissues and conditions were compared using the publicly available data from PeatMOSS (Fernandez-Pozo et al., 2020).

For comparison of conservation degree of COX11, the protein sequences from cattle (*Bos taurus)*, UniProt ID A3KMZ6; yeast (*Saccharomyces cerevisiae*), UniProt ID P19516; and *P. patens*, UniProt ID A0A2K1J6Q0_PHYPA were retrieved and then aligned using Clustal Omega v. 1.2.4 (Sievers et al., 2011).

### Western blot analysis

Protein extraction and immunoblotting was performed as described previously (Mellon et al., 2021). Samples were loaded so that the same amount of chlorophylls was present. For chlorophyll quantification, 2 mL of pure protein extract were diluted in 68 μL of acetone 80 % in 0.5 mL tubes. The tubes were vortexed briefly and centrifuged at 13,000 g for 5 min at room temperature. Proteins precipitated forming a pellet and chlorophylls remained in solution. The supernatant was transferred to clean 0.5 mL tubes. The absorption spectrum (from 750 to 600 nm) of the supernatant was measured using a Cary 100 UV-Vis spectrophotometer (Agilent Technologies). The chlorophylls concentration of the pure protein extract, [*Chl*], was calculated applying the following formula, where OD stands for optical density (Porra et al., 1989):

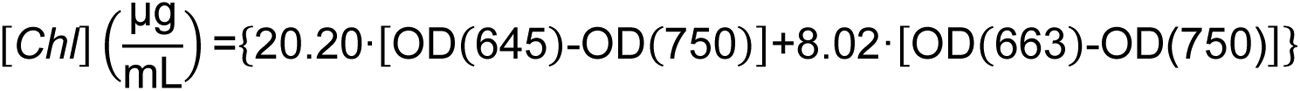

After quantification, the pure protein extracts were incubated 1 min at 100 °C.

The list of antibodies used is attached as Supplementary Table 2. For all the antibodies except for the ß-subunit of complex V, total protein extracts were resolved by SDS-PAGE. For the ß-subunit of complex V, crude mitochondria extracts were separated by Urea-PAGE.

### Crude membrane isolation, blue native protein electrophoresis (BN-PAGE) and complex IV activity staining

Crude membrane extracts were prepared on ice and using chilled tubes and reagents to avoid protein denaturation following a protocol adapted after Pineau et al., 2008. Approximately 300 mg of fresh or frozen (-80 °C) protonema grown on PpNO3 for 10 days were homogenised using a Potter-Elvehjem glass tissue grinder in 2 mL of MOPS-KOH 75mM, pH 7.6; sucrose 0.6 M; EDTA 4 mM; polyvinylpyrrolidone-40 0.2 %; cysteine 8 mM and bovine serum albumin 0.2 %. The homogenate was transferred to a 15 mL conical tube lined with a square of miracloth with 20 μm pores to filter unbroken cells and tissue debris. The filtered homogenate was then transferred to a 2 mL tube and centrifuged for 4 min at 1,300 g to pull down the cellular debris. The supernatant was collected in a clean 2 mL tube and centrifuged for 20 min at 21,470 g to pellet the thylakoid and mitochondrial membranes. The supernatant was discarded, and the pellet was resuspended in 200 μL of MOPS-KOH, pH 7.2 and sucrose 0.3 M. Total protein content was determined by the bicinchoninic acid (BCA) assay (He, 2011).

A 4-12 % running acrylamide gradient polyacrylamide gel was freshly prepared, with a stacking 4 % gel. Protein solubilization protocol was adapted after Pineau et al., 2008. 20 µL of digitonin 4 % (w/v) prepared in ACA buffer (1.5 M aminocaproic acid, 0.1 M Bis-Tris-HCl pH 7.0, 2 mM EDTA) were added to the tubes containing 20 µL of crude extracts in ACA buffer that corresponded to 50 µg of proteins, reaching 2 % digitonin in ACA buffer. Each tube was incubated on ice for 20 min and centrifuged at 4 °C, 22,000g for 8 min. The supernatant containing solubilized protein complexes was transferred to a clean tube, supplemented with 4 µL of Coomassie Blue 5 % solution (20 mM Bis-Tris, 0.5 M aminocaproic acid, Coomassie Blue G-250 5 % (w/v)) and loaded to the gel.

Sample loading and gel running were performed in a cold chamber at 4 °C according to an adapted version of the protocol described by Järvi et al., 2011. The electrode assembly with the mounted gel was filled with both cathode buffer (15 mM Bis-Tris-HCl, pH 7.0; 50 mM tricine) containing Coomassie Blue G-250 0.02 % (w/v) and anode buffer (50 mM Bis-Tris-HCl, pH 7.0). The wells were loaded with the solubilised samples using a Hamilton syringe. The gel was run at 75 V for 30 minutes. Then, the cathode buffer was replaced with fresh cathode buffer without Coomassie Blue and the gel was run at 100 V for 30 min, at 125 V for 30 min, at 150 V for 60 min, at 175 V for 30 min and at 200 V for 60 min. The total running time was about 4 hours. After running, the gel was kept at 4 °C until used for in-gel activity staining.

After running, in-gel activity staining was done as described in (Sabar et al., 2005). The gel was incubated for 1 h in complex IV buffer (50 mM potassium phosphate buffer pH 7.4, 75 mg/mL sucrose). Then, the buffer was replaced with complex IV staining solution [1 mg/mL DAB (3,3′-Diaminobenzidine tetrahydrochloride hydrate; Merck D5637), 1 mg/mL cytochrome c (Merck C7752)] in complex IV buffer). The gel was incubated in the staining solution for 24 h at room temperature before image acquisition.

### In-gel amylolytic activity staining

For in-gel amylolytic activity, proteins were extracted 1:2 (fresh weight: volume) in 100 mM MOPS pH 7.2, 1 mM EDTA, 10% glycerol, 5 mM DTT, and 1 mM phenylmethylsulfonyl fluoride using a pestle. Proteins were quantified by the BCA assay (He, 2011). For each sample, 10 µg were mixed to native loading buffer (60 mM Tris-HCl pH 6.8; 10 % glycerol; 0.025 % bromophenol blue) and loaded into a 7.5 % acrylamide gel containing 0.1 % solubilized amylopectin from potato starch (Merck). Gels were run at 120 V for 4 h at 4 °C and then incubated in incubation buffer (100 mM Tris-HCl pH 7; 1 mM MgCl_2_; 1 mM CaCl_2_, 2 mM DTT) for 15 min, the buffer was changed with new buffer and incubation left overnight at room temperature or at 37 °C for 4 h. After the incubation, gels were washed with bidistilled water and stained with Lugol’s solution (0.33 % I_2_, 0.66 % KI).

### Growth test

2-mm disks of protonema of *P. patens* grown for 10 days in long day conditions were distributed in 92 mm Petri dishes filled with either solid PpNO3, PpNH4 or amino acid-enriched media. A cellophane filter was not added as routinary done for other experiments. Each Petri dish contained 10 plant disks and at least two technical replicates per genotype were included. The space distribution of the disks on the plate was homogenous and the relative position of each disk was aleatory, but the same loading scheme was followed for the different conditions to be compared. Colony size was measured periodically. Photographs of the plates were acquired using a smartphone. Images were processed with Fiji (Schindelin et al., 2012) to remove the plate background using the “Threshold Colour” plugin and/or the “Threshold…” utility after image conversion to 8-bit. Scale was set using the diameter of the Petri dish (92 mm) and area in mm^2^ was measured for each colony.

The effect of photoperiod was assessed by growing identical plates under either long day or continuous illumination (24 h light). The effect of high CO_2_ was evaluated by keeping the Petri dishes in a growth chamber with controlled atmosphere of 1 % CO_2_ in continuous illumination and 22 °C. The effect of amino acid addition was tested by performing the growth test on solid PpNO_3_ medium containing additional 3 mM of each of the twenty primary amino acids except for tyrosine, which could not be dissolved without strongly acidifying the medium. The final concentration of 3 mM was chosen according to previous experiments done in Arabidopsis (Forsum et al., 2008).

### Measurement of oxygen consumption and evolution

Measurements of oxygen consumption (respirometry) and oxygen evolution were performed on pieces of intact protonema from 10-day old plants grown on PpNO3 and dark adapted for 40 minutes before the experiments. Measurements were performed using a test version of the NextGen-O2k and the PhotoBiology (PB)-Module (Oroboros Instruments, Innsbruck) with the software DatLab 7.4.0.4 (Went et al., 2021). The PB light source contained a blue OSLON® LED (emitting wavelength range 439-457 nm with the peak at 451 nm) attached to the window of the NextGen-O2k chamber. The oxygen concentration was assessed in 2-mL measuring chambers at 22 °C with a 2-seconds frequency and samples were magnetically stirred at 750 rpm.

To avoid disruption of the moss samples during the measurement, we used a sample holder developed by Oroboros Instruments, Innsbruck (Schmitt & Gnaiger, 2022). At first, we filled the measuring chambers with a volume slightly higher than 2 mL of fresh and sterile PpNO3 medium containing 10 mM NaHCO_3_ (to avoid carbon limitation during photosynthetic measurements), we inserted the sample holder and closed the chamber removing excess volume to ensure that precise 2 mL were inside. Then, we moved the stopper to the open position and let the system equilibrate for few minutes with the stirring on. This served both to bring the medium to the experimental temperature and to equilibrate the oxygen concentration of the medium to the atmospheric oxygen values. We opened the chamber, added a piece of protonema of approximately 1 cm^2^ on top of the sample holder and closed the chamber. This operation was done minimizing the exposure of moss samples to light. Oxygen concentration in the chamber was monitored for 10 minutes at dark to assess the rate of dark respiration, and we proceeded either with the quantification of photosynthesis or with the assessment of the effect of respiratory inhibitors.

For quantifying oxygen evolution, after stabilization of the respiration signal, blue light was turned on at 500 μmol photons m^-2^ s^-1^, which was well above the saturating levels for *P. patens*. We kept the sample under this illumination for 10 minutes to achieve the stabilization of the oxygen evolution rate. The values of oxygen evolution rate reported in this work correspond to the median of 40-50 points in the stable region of oxygen flux. The plot of a representative experiment is shown as Supplementary Figure 8.

For quantifying the effect of inhibitors on dark respiration, after stabilization of the respiration signal, inhibitors were added sequentially to the chamber. The plot of a representative experiment is shown as Supplementary Figure 6. Inhibitors were added through the stopper using Hamilton syringes, therefore not interrupting the measurements. For titrations using KCN or Antimycin A, we added to the chamber 4 µL of a stock 500 mM; for titrations using SHAM or n-propylgallate, we added to the chamber 8 µL of a stock 250 mM. In all cases the final concentration of the inhibitor was 1 mM.

After each experiment, the moss sample was recovered from the chamber and used for chlorophyll quantification as described earlier in this section. The measure of oxygen consumption and evolution was therefore normalized to the amount of chlorophylls in the sample.

### Measurement of CO2 consumption and evolution

Quantification of CO_2_ exchange was done using a homemade atmosphere simulation chamber initially developed for its use with cyanobacteria as described in (Battistuzzi et al., 2020, 2023). A single 92-mm Petri dish containing 10-day old protonema was introduced inside of the sealed chamber and the net CO_2_ consumption rate was measured under illumination.

### Spectroscopic analyses

*In vivo* Chl fluorescence analyses were performed as described previously (Mellon et al., 2021).

### Transmission electron microscopy (TEM)

10 days old plants grown under long day were kept at dark for circa 20 h before fixation, to avoid overaccumulation of starch. Small pieces of samples (about 2-3 mm^3^) were fixed with 3.0 % glutaraldehyde (EMS 16220) plus 1 % paraformaldehyde (Merck P6148) plus 0.5 % tannic acid in 0.05 M sodium cacodylate buffer pH 7.4 for 2 h at RT. Subsequently the samples were postfixed with 1 % in 0.1 M sodium cacodylate buffer for 1 hour at 4 °C. After three water washes, samples were dehydrated in a graded ethanol series and embedded in an epoxy resin (Sigma-Aldrich 46345). Ultrathin sections (60-70 nm) were obtained with a Leica Ultracut EM UC7 ultramicrotome, counterstained with uranyl acetate and lead citrate and viewed with a Tecnai G2 (FEI) transmission electron microscope operating at 100 kV. Images were captured with a Veleta (Olympus Soft Imaging System) digital camera.

### Sample harvesting for systems analysis

We optimized the following system for growing moss samples for systems level analyses, i.e. RNA sequencing and untargeted metabolomics (Supplementary Figure 13). Protonema grown for one week on solid PpNH4 medium under continuous illumination was disrupted using a T 25 UltraTurrax (IKA) and used for the inoculation of a 100 mL Erlenmeyer flask with 20 mL of liquid PpNH4. After one week of growth under continuous illumination, the plant was harvested, disrupted and used for the inoculation of a 250 mL Erlenmeyer flask with 50 mL of liquid PpNO3. After one week of growth under continuous illumination, the plant was harvested, disrupted and used for the inoculation of a 500 mL Erlenmeyer flask with 100 mL of liquid PpNO3. After one week of growth, the plant material was distributed into a layer of miracloth arranged on top of a plastic cylinder, inside of a Magenta™ box filled with fresh liquid PpNO3. The plant was in contact with the medium through the filter, allowing growth under this “hydroponics” system. Plants in closed Magenta™ boxes were grown for 15 days under a short day regime (light:dark 12h:12h). The day of harvesting, plants were immediately snap frozen at the corresponding zeitgeber time (ZT0, ZT2, ZT6). For plants harvested at ZT0, Magenta™ boxes were enclosed in aluminium foil at the beginning of the night period (ZT12) and samples were snap frozen at ZT0 in a dark room illuminated with dim, green light. For plants harvested at ZT2 or ZT6, plants were snap frozen directly in the growth chamber, avoiding shading the plant until it had been frozen. Frozen samples were kept at -80 °C until used.

### RNA sequencing, differential gene expression analysis and pathway enrichment analysis

Frozen samples were powdered using a cold mortar and pestle, in presence of liquid nitrogen. Approximately 150 mg of powder were used for RNA extraction using the RNeasy Plant Mini Kit, ref. 74904 (Qiagen). RNA was used to build cDNA libraries using the kit QuantSeq 3’ mRNA-Seq Library Prep Kit FWD (Lexogen), which exploits oligo-d(T) primers therefore enriching the cDNA library in cDNA representative of mRNA, but not ribosomal RNA or other types of RNA. Libraries were then sequenced with a depth of 5 million of reads using a NextSeq 500 System (Illumina).

The reference genome (Physcomitrium patens v3.3) from the JGI Plant Gene Atlas (Lang et al., 2018) was indexed using Bowtie 2 (v2.2.5) (Langmead & Salzberg, 2012). Reads were then aligned using Bowtie 2 and bam files were generated with SAMtools (v 1.6) (Danecek et al., 2021). HTSeq-count (v 2.0.2) (Putri et al., 2022) was used to align the mapped reads to the genes provided in the annotation file (Ppatens_318_v3.3.gene.gff).

Differential expression analysis was done through the tool iDEP (v 0.96) (Ge et al., 2018). First, a low filter was applied to the gene list to keep only genes with at least 50 reads in at least one sample, which were 16,101. Then, their expression values of were log_2_ transformed, applying the formula x’ = log_2_(x+1). Differential expression was analysed using the integrated *limma* package (Ritchie et al., 2015), setting the false discovery rate (FDR) cutoff at 0.05 and the fold-change cutoff at 2. The lists of differentially expressed genes (DEGs) at each condition are given as Supplementary Dataset 1. These lists were used for pathway enrichment analysis with iDEP using all the annotation databases available for *P. patens* in iDEP, which were based in Gene Orthology (GO), or annotated protein domains coming from different sources (UniProt, InterPro, Pfam, SMART). The lists and composition of enriched pathways at different conditions are given as Supplementary Dataset 2.

The list of genes encoding for ascorbate peroxidase was retrieved from three different sources. Some of them had alternative names in different publications. The summarized information is included as Supplementary Table 3. The list of genes encoding for superoxide dismutases and glutathione reductases were retrieved from Higashi et al., 2013 and L. Xu et al., 2013, respectively. For glutathione reductase, the five genes defined as H_2_O_2_-responsive by Y. J. Liu et al., 2013 were included.

### Untargeted metabolomics and integrated pathway analysis

To a 1.5 mL tube containing 50-100 mg of powdered, frozen moss samples, we added 350 µL of extraction solution (chloroform:methanol 10:4.28), vortexed and incubated at -20 °C for 1 h. Then we added 560 µL of internal standard stock solution, kept on ice for 5 min with frequent vortexing, and centrifuged for 2 min at 20,000 g. We then collected the supernatant into fresh 2 mL tubes and performed a second extraction of the remaining organic phase by adding 260 µL of H_2_O, followed by incubating on ice for 5 min with frequent vortexing and centrifugation for 2 min at 20,000 g. The second supernatant was added to the previous one. The extracts were kept at -80 °C until analysis by two different methods: GC-MS (according to Shim et al., 2020) and IC-MS (according to Curien et al., 2021). Since we produced 2 or 3 biological replicas for each genotype for each condition, we combined the replicas from the two independent *cox11* lines into a broader group that contained 4 to 6 replicas per each condition. This increases the power of statistical analyses. The abundance of metabolites was normalized to the internal standard and to the dry weight.

We then used the data as input values for the publicly available tool MetaboAnalyst (Xia & Wishart, 2011), and used the graphics user interface to generate volcano plots and compare metabolite levels between mutant and WT at the different zeitgeber times. As an output we obtained the log-fold change values and *p* values of every single comparison. To elaborate lists of metabolites that were significantly accumulated or depleted at a given condition, we established the significance threshold at *p*<0.1. Heatmap in Figure 6 was obtained using the platform SRPlot (Tang et al., 2023)

Adenylate Energy Charge (AEC) was calculated as (ATP + ½ ADP) / (ATP + ADP + AMP) according to Tyutereva et al., 2022, based on the relative concentration of ATP, ADP and AMP obtained by IC-MS after normalization to the internal standard thio-ATP.

Integrated pathway analysis was done by visualizing transcriptomics and metabolomics data with Pathway Tools (Karp et al., 2021) using the MossCyc v8.0.2 database from the Plant Metabolic Network (Hawkins et al., 2021). Metabolic diagrams in Figure 6B were manually produced.

### Quantification of starch

Quantification of starch was done on total extracts of 10 days-old *P. patens* samples following the protocol described in A. M. Smith & Zeeman, 2006.

## Accession numbers

Accession numbers of gene sequences: Cox11, Pp3c16_1230; Actin7, Pp3c3_33440. Accession numbers of specific pathways are included in supplemental tables 3-5.

## Supplementary Data

**Supplementary Figure 1.**
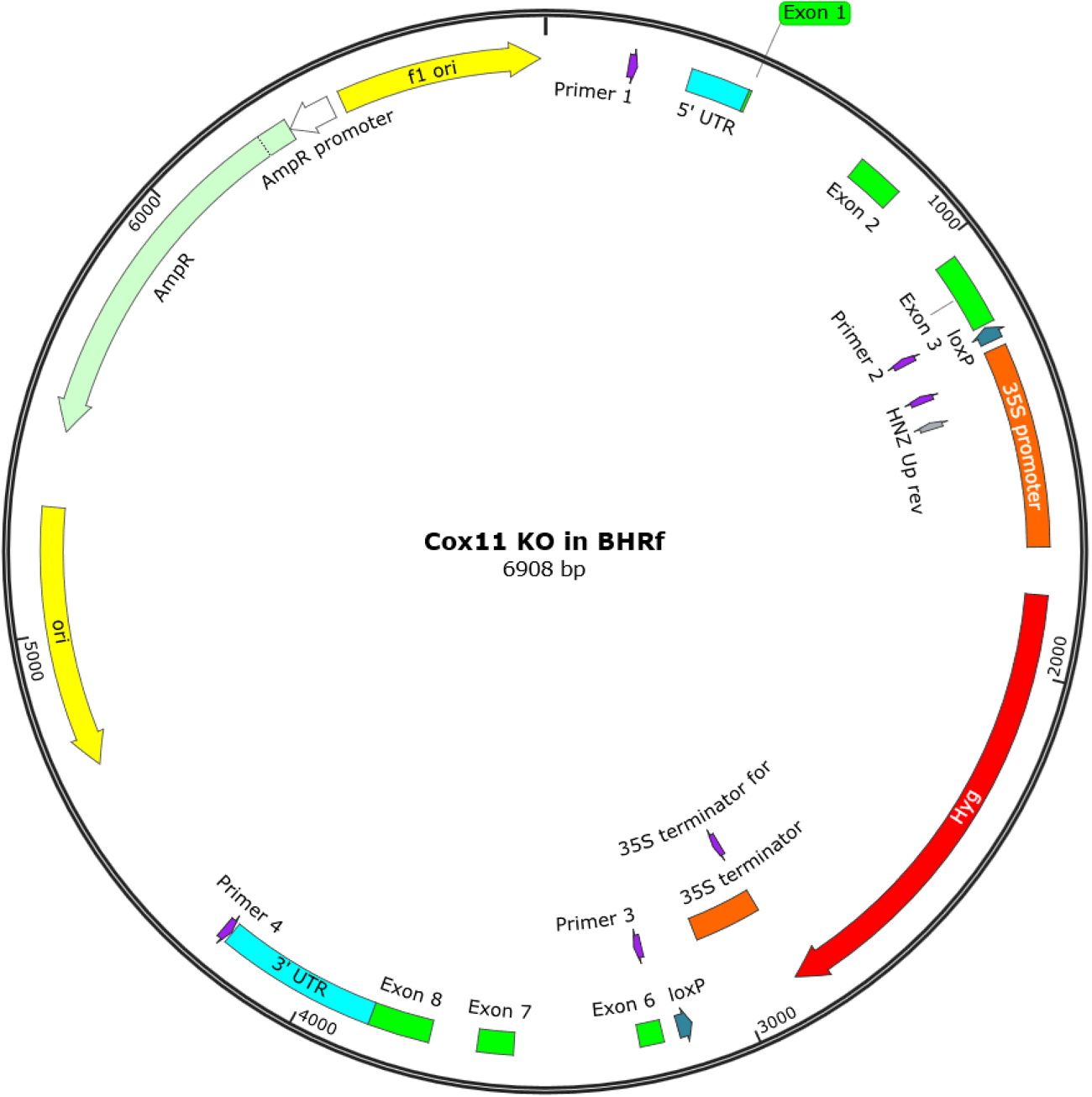
Map of the construct used for knocking out *COX11*. Primer sequences are included in Supplementary Table 1.

**Supplementary Figure 2.**
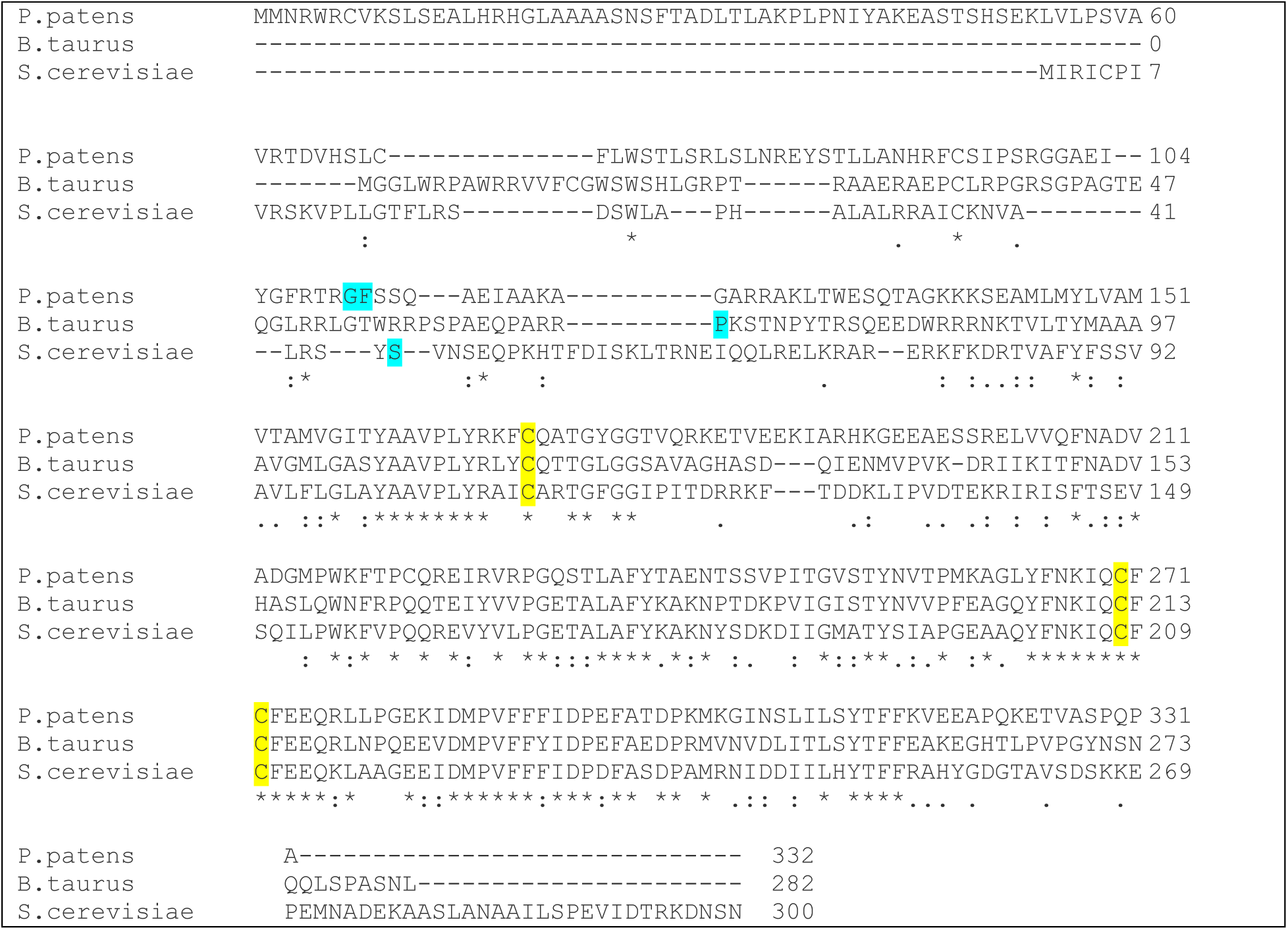
Multiple sequence alignment of protein Cox11 of *P. patens* and their homologs in yeast (*Saccharomyces cerevisiae*) and cattle (*Bos taurus*). The multiple sequence alignment was performed using the tool *Clustal Omega (1.2.4).* Letters with cyan background indicate the cleavage sites predicted by the tool *MitoFates* (Fukasawa et al., 2015). Letters with yellow background are the conserved Cys residues involved in copper binding as identified by Carr et al., 2002.

**Supplementary Figure 3.**
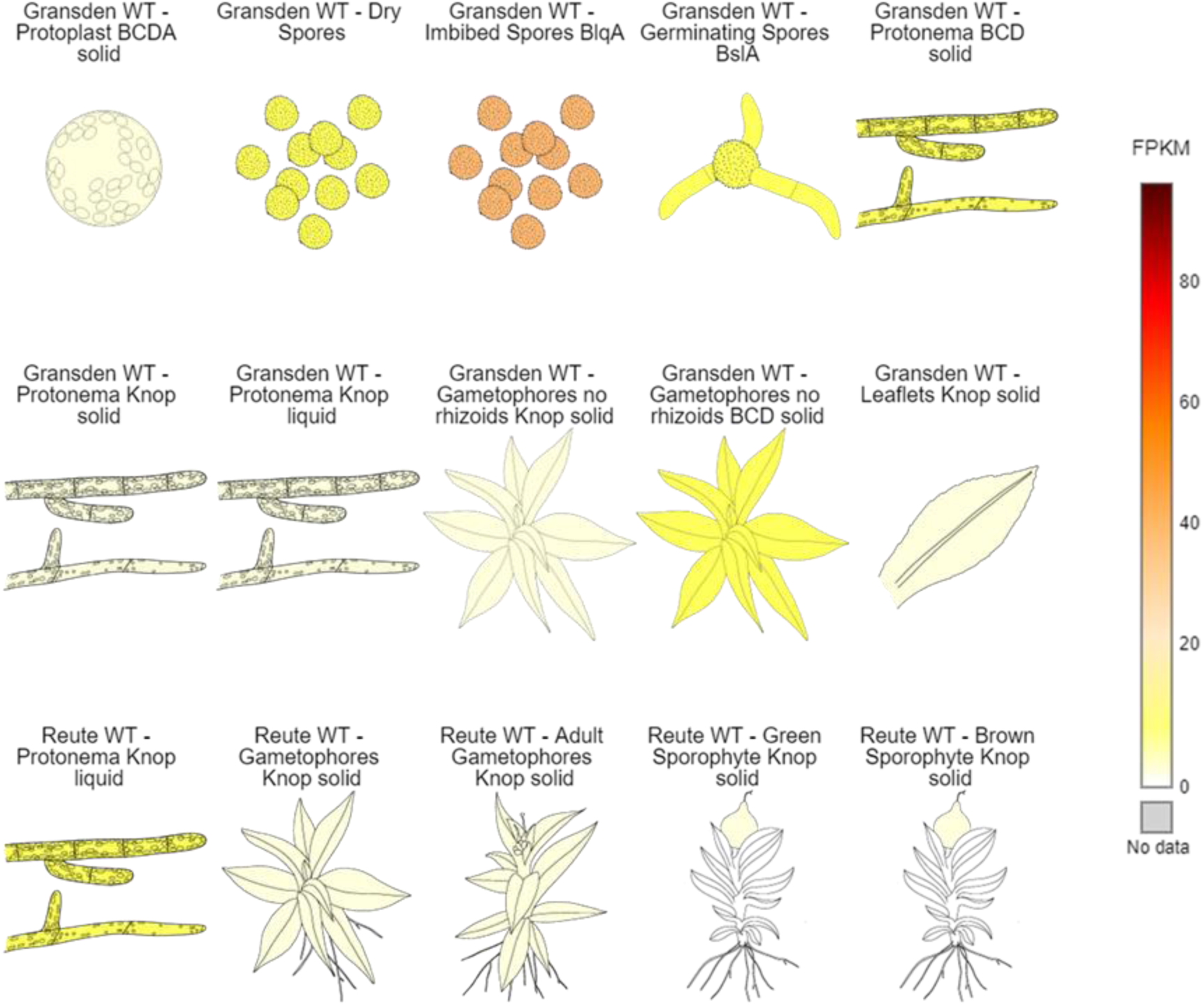
Expression levels of *COX11* in different developmental stages. Note that the FPKM are relatively low for all stages, with imbibed spores presenting the highest expression. Data and figure were retrieved from the PeatMOSS Gene Atlas Database (Fernandez-Pozo et al., 2020).

**Supplementary Figure 4.**
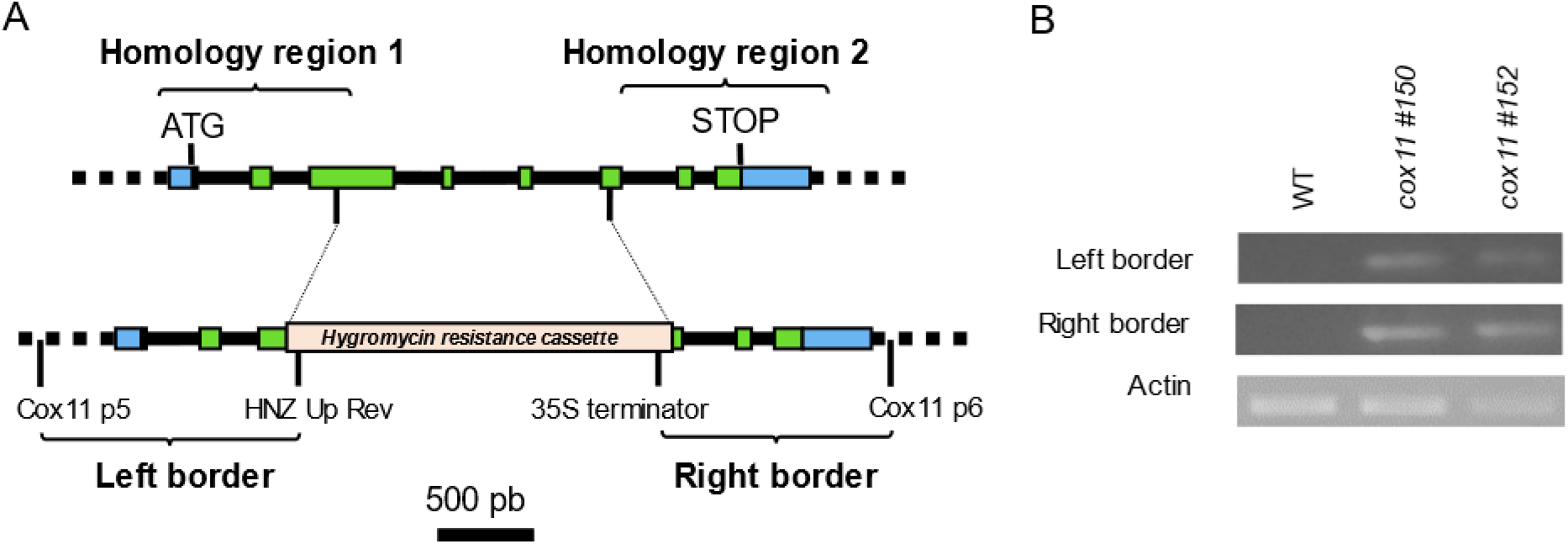
Visual summary of the disruption of the *COX11* locus used for producing the knockout mutant lines. The gene map (A, top) and the corresponding region after construct insertion through homologous recombination (A, bottom) are shown. Green portions represent the exons, blue portions represent the 5’ and 3’ untranslated regions (UTR). The positions of the first and last codon of each coding region are marked with ATG and STOP, respectively. Integration of the antibiotic resistance cassette occurs through recombination of the respective homology regions. The two regions used for validation of their identity through PCR analysis using the given primers are marked as Left Border (LB) and Right Border (RB), and the corresponding bands are shown in B. Primer sequences are included in Supplementary Table 1.

**Supplementary Figure 5.**
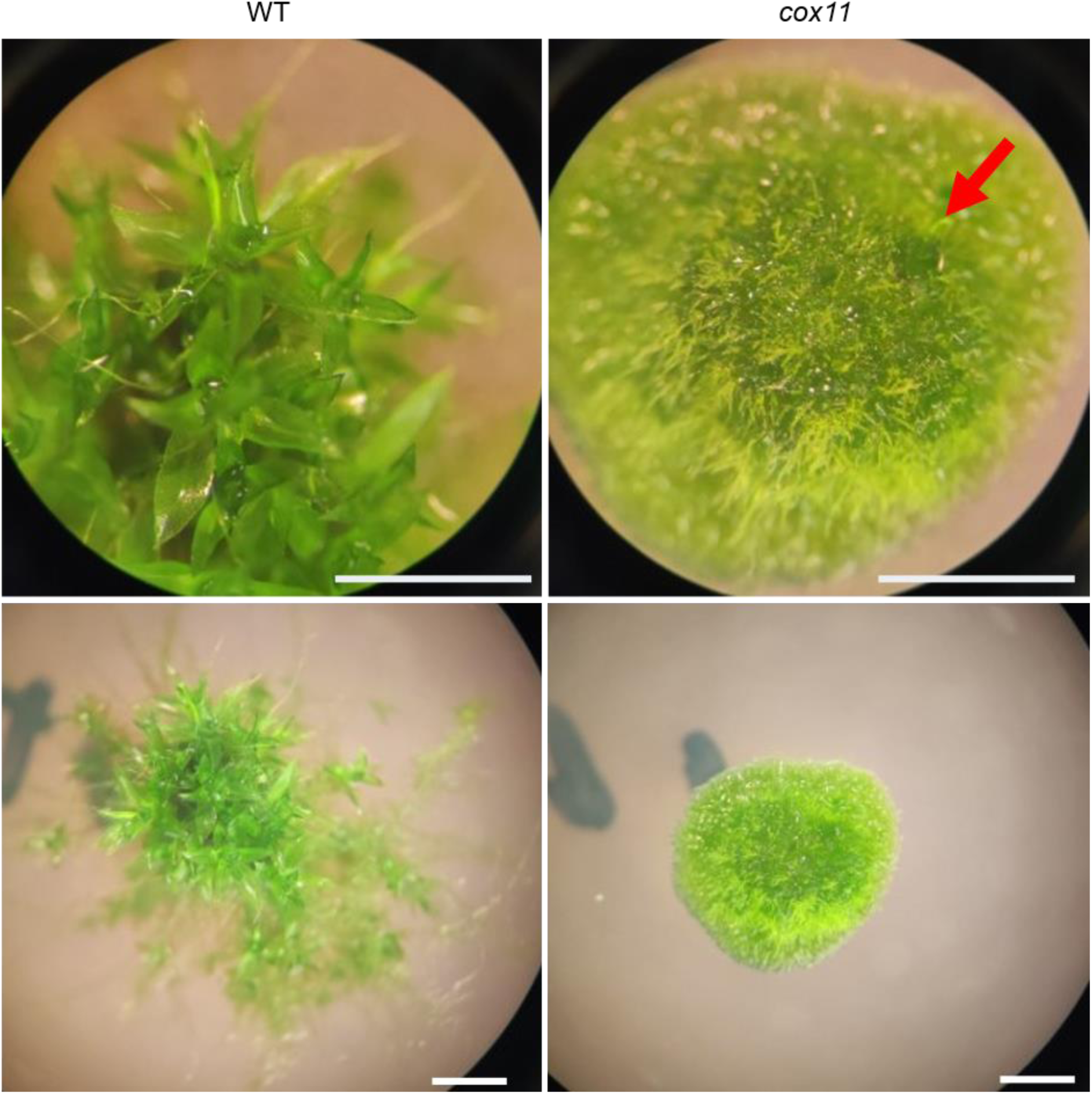
Differential development of gametophores in *cox11*. Images of plant colonies after 21 days of growth, at two different magnifications. A developing gametophore is marked with a red arrow in *cox11*. Scale bars are 2 mm.

**Supplementary Figure 6.**
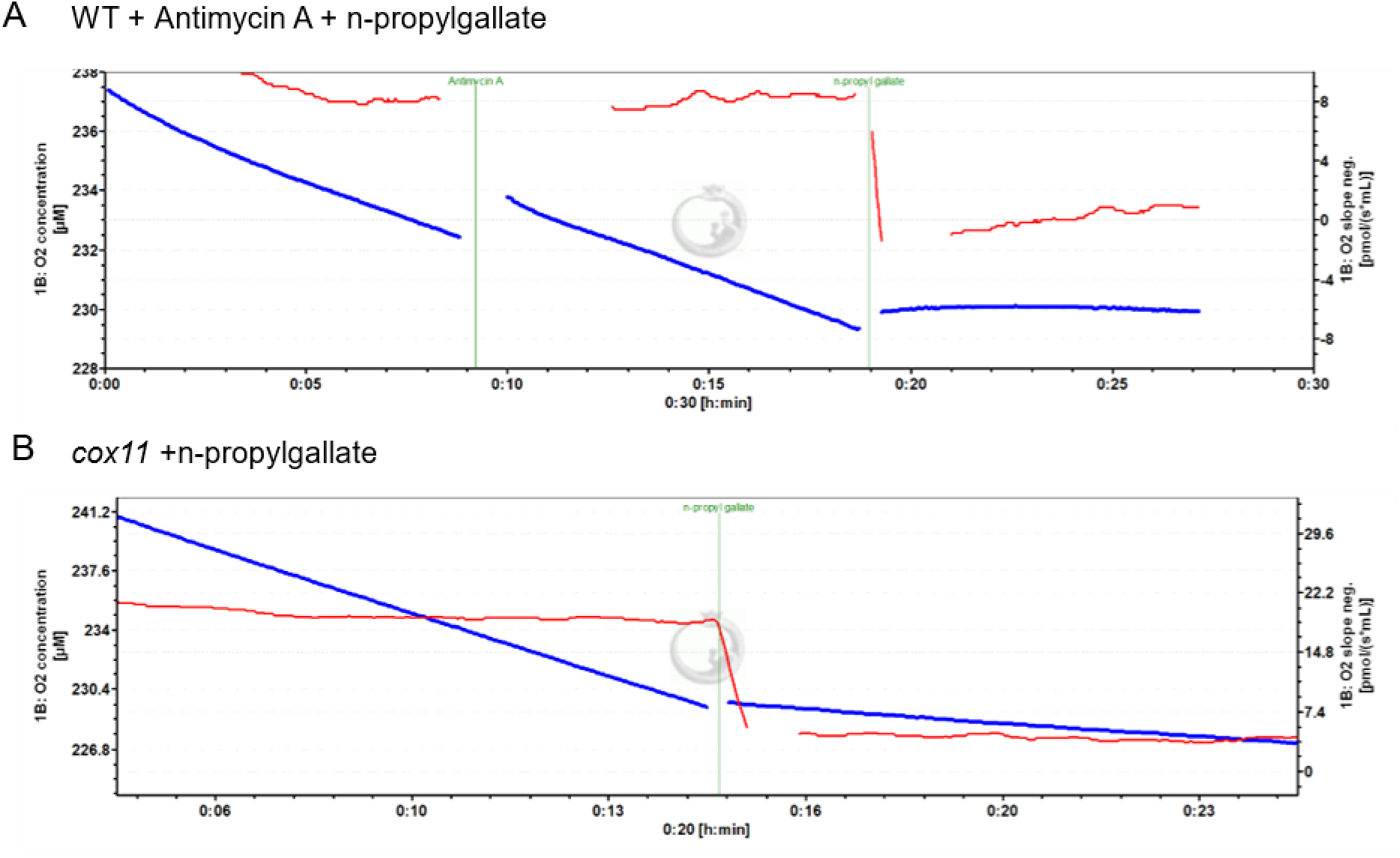
Effect of antimycin and n-propylgallate on dark respiration of intact protonema. Two representative tracks of experiments of respirometry performed on WT (A) or *cox11* (B) are shown. The blue plot (left axis) shows O2 concentration, while the red plot (right axis) shows O2 flux. On WT, the addition of Antimycin A did not have an effect in O2 consumption rate, but n-propylgallate abolished it (A). On *cox11*, n-propylgallate alone was sufficient to block all O2 consumption (B). Images are screenshots of the DatLab software.

**Supplementary Figure 7.**
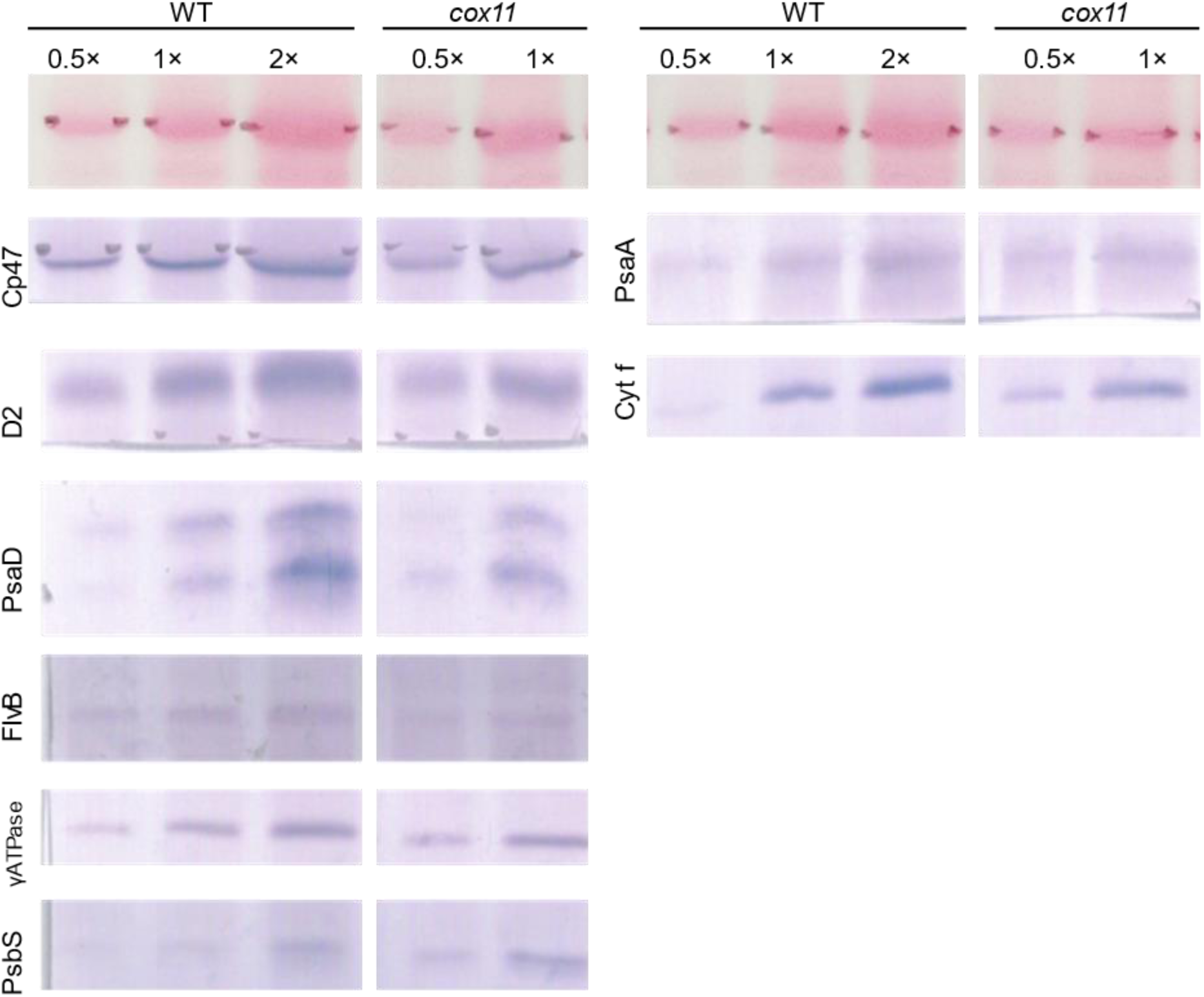
Immunoblotting against subunits of the photosynthetic machinery. No major alterations in photosynthetic components were detected. 1× corresponds to 2 µg of loaded chlorophylls.

**Supplementary Figure 8.**
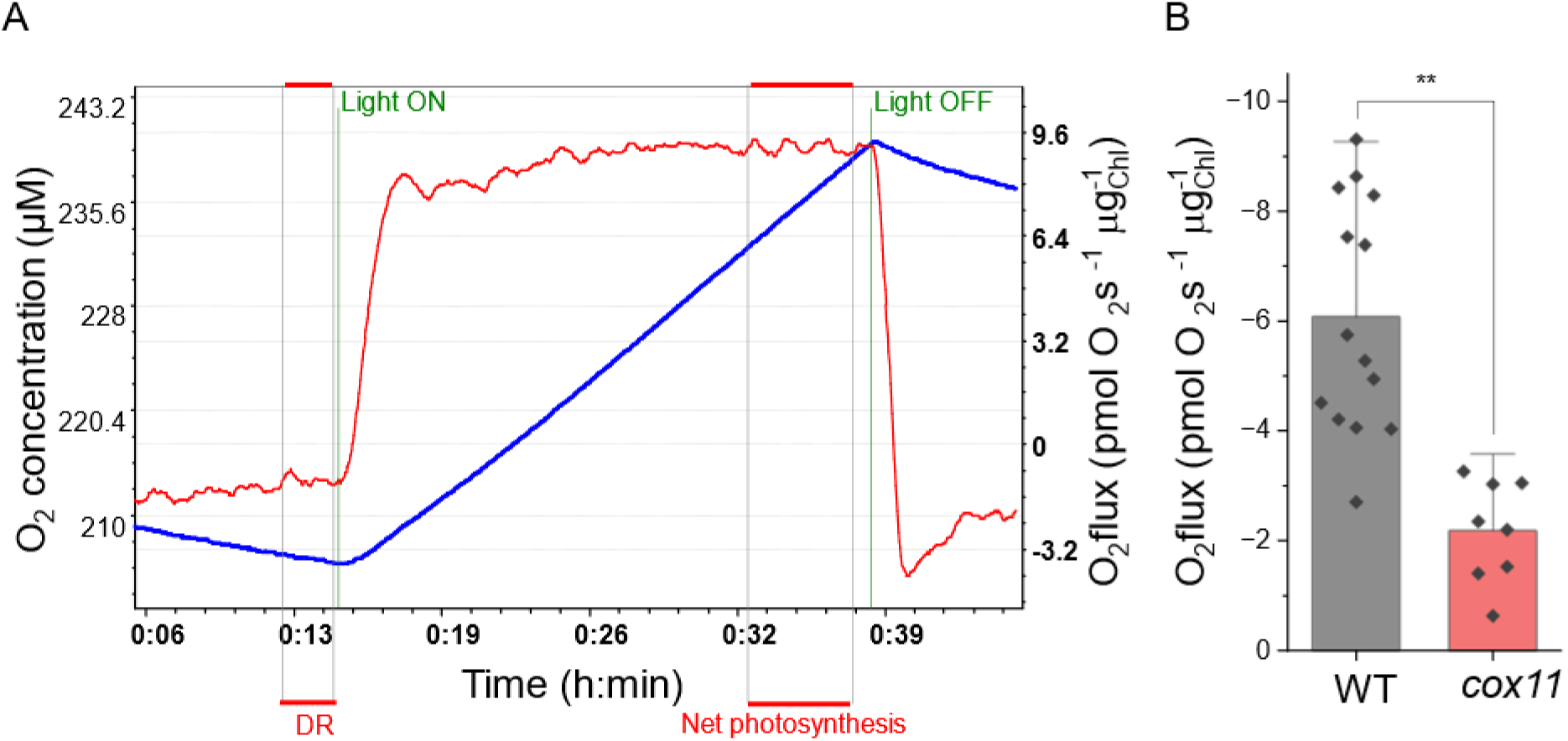
Net photosynthesis measurement and values on intact protonema. (A) Representative track of one experiment of respirometry. The blue plot (left axis) shows O2 concentration, while the red plot (right axis) shows O2 flux. After quantification of dark respiration (DR), saturating light was turned on. When the O2 flux was stable (red plot), the net photosynthesis was measured as the median of the shown stable region. (B) Values of net photosynthesis from WT and *cox11*. Values in *cox11* were obtained from two independent lines. Statistics: (**) p<0.01.

**Supplementary Figure 9.**
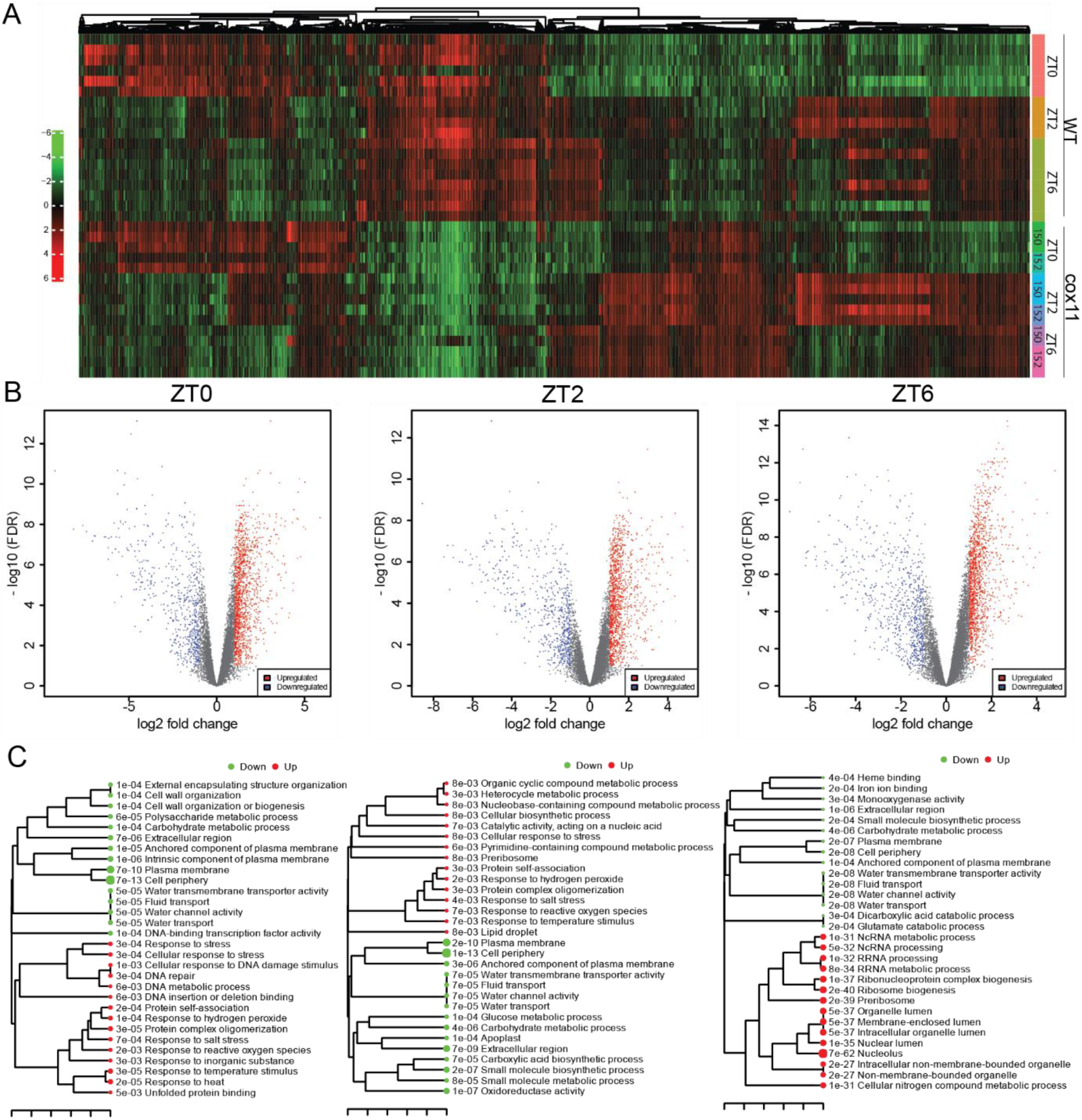
RNA-seq data overview and enriched pathways segregated for the three zeitgeber times tested. (A) Hierarchical clustering of the 2,000 top genes, showing Pearson distance and average linkage. For each zeitgeber time, a volcano plot showing data distribution and the number of significant DEGs (B) and a tree of the most significant enriched pathways, where pathways with many shared genes are clustered together, and bigger dots indicate more significant p-values (C), are reported.

**Supplementary Figure 10.**
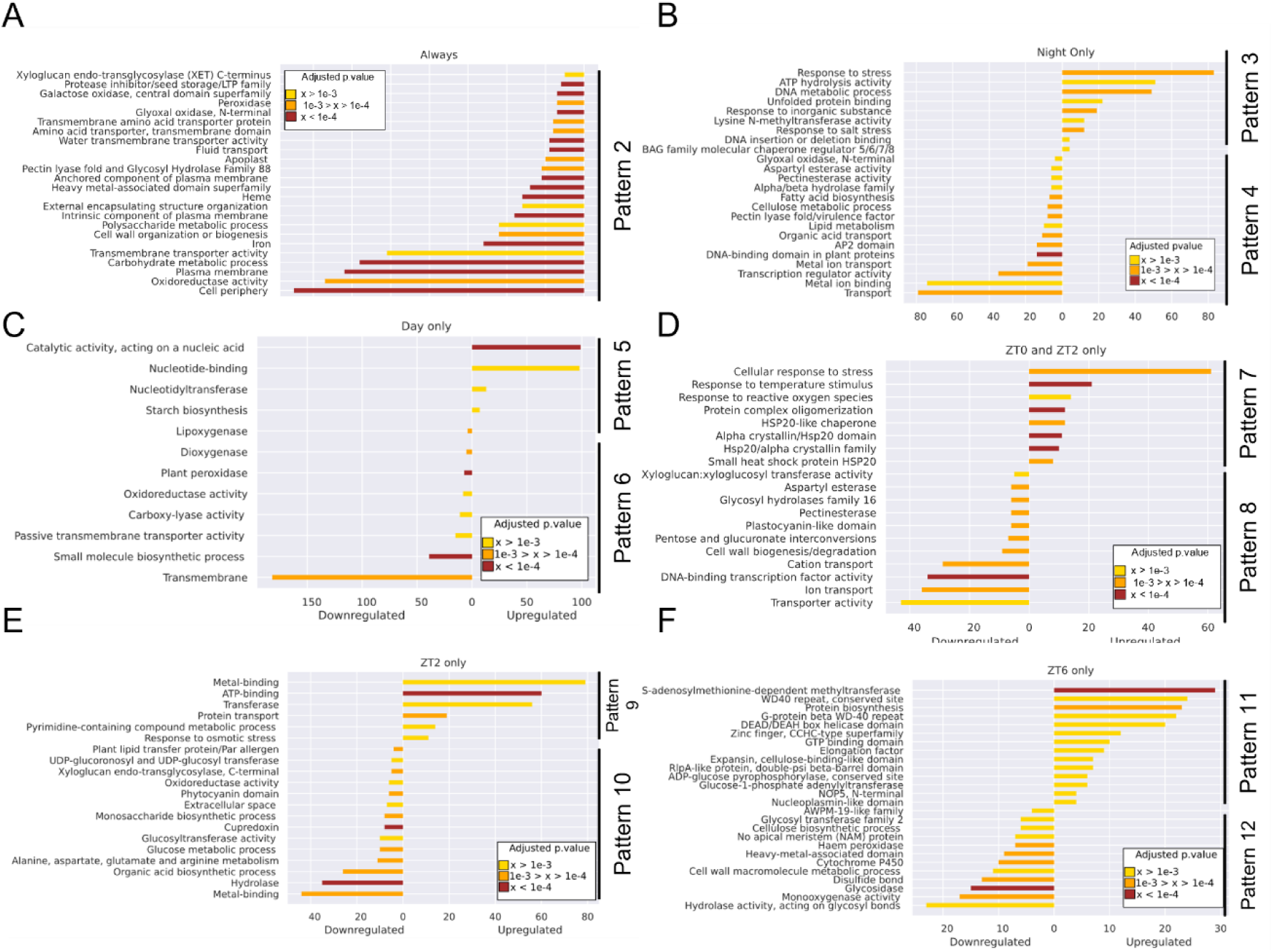
Bar plots with the most significant pathways with differential expression in *cox11.* Different groups are distinguished following the pattern classification defined in Figure 5. Length of bars represents numbers of genes included, while colours represent adjusted p-values of the enriched pathway. Detailed information and composition of these pathways can be found at **Supplementary Dataset 2**.

**Supplementary Figure 11.**
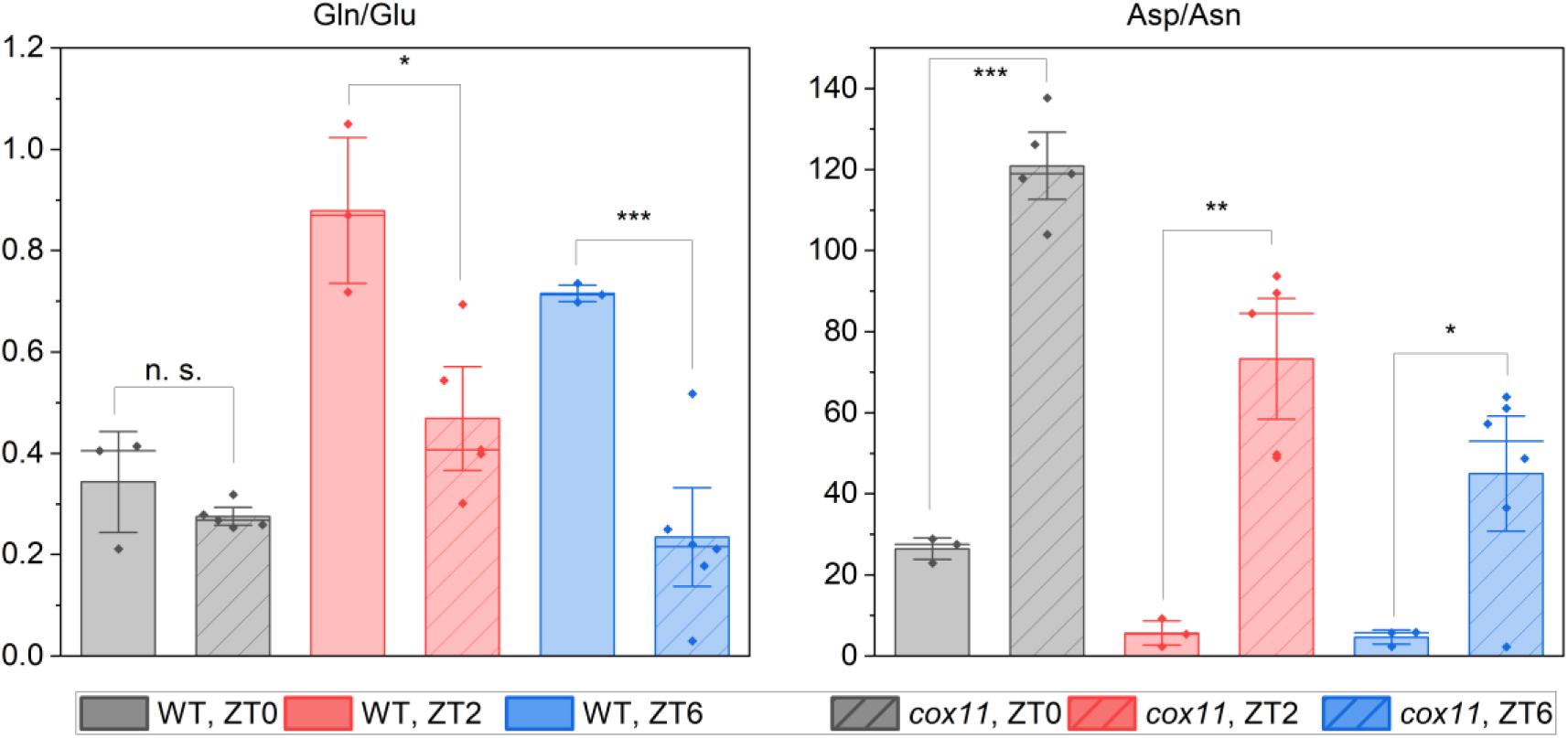
Ratios comparing relative abundances of glutamate/glutamine and aspartate/asparagine. Statistics: two-sample t-test, (***) p<0.001; (**) p<0.01; (*) p>0.05; (n. s.) not significant.

**Supplementary Figure 12.**
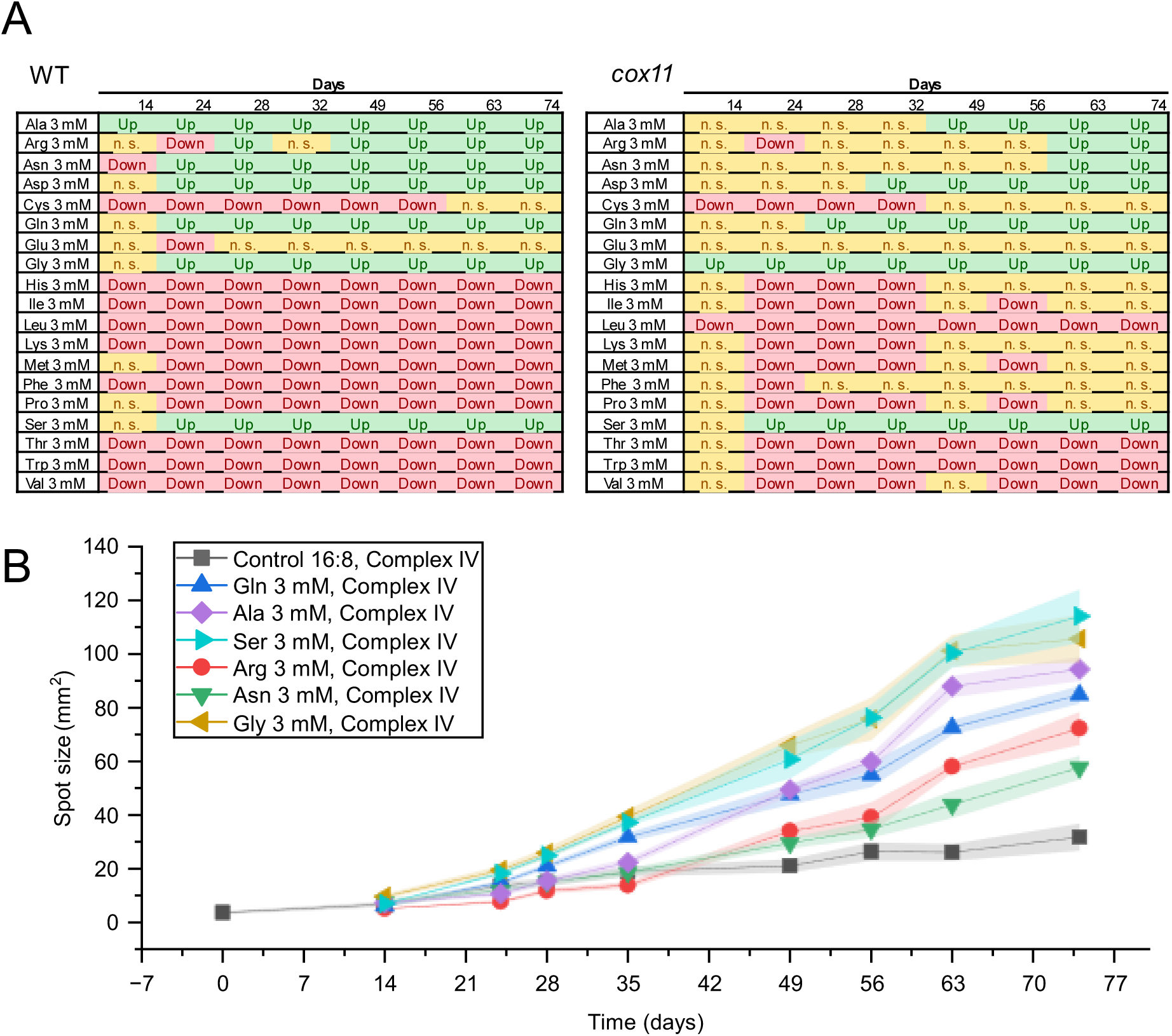
Effect of different amino acids on *cox11* growth. (A) Time-resolved effect on growth of addition of the different amino acids. The size of the spots was quantified at each given timed and compared with the control by multiple two-way ANOVA; means comparison was done through Fisher LSD (least significant difference) test. For significant comparisons (*p*<0.05), the direction of growth is shown (Up, Down). Not significant comparisons are depicted as “n. s.”. (B) Area of the spots of *cox11* plants grown on PpNO3 supplemented with amino acids. Only the amino acids that caused an improvement of growth in *cox11* are included.

**Supplementary Figure 13.**
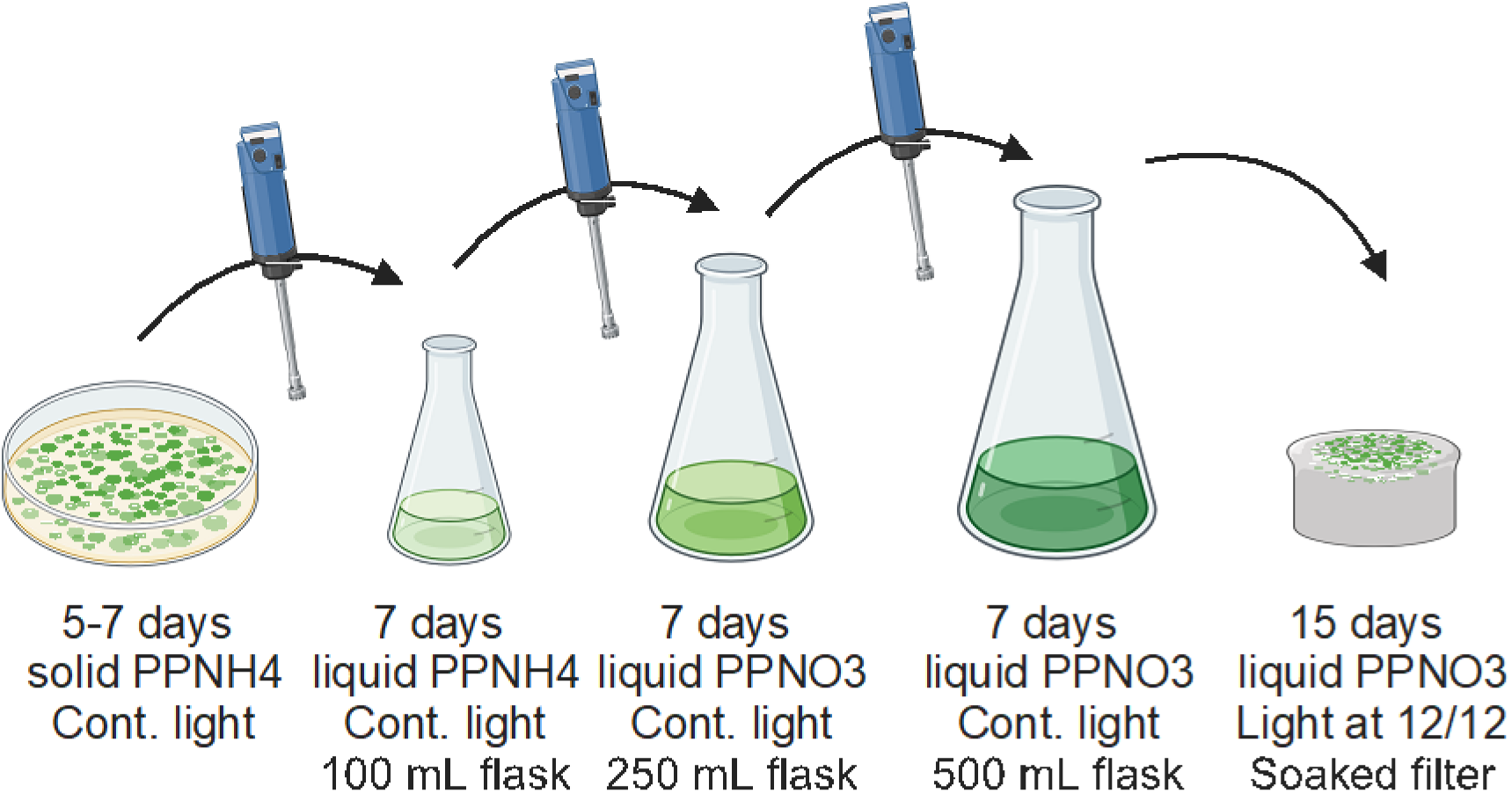
Protocol for amplifying moss material for transcriptomics and metabolomics. Created with BioRender.com.

**Supplementary Table 1.**
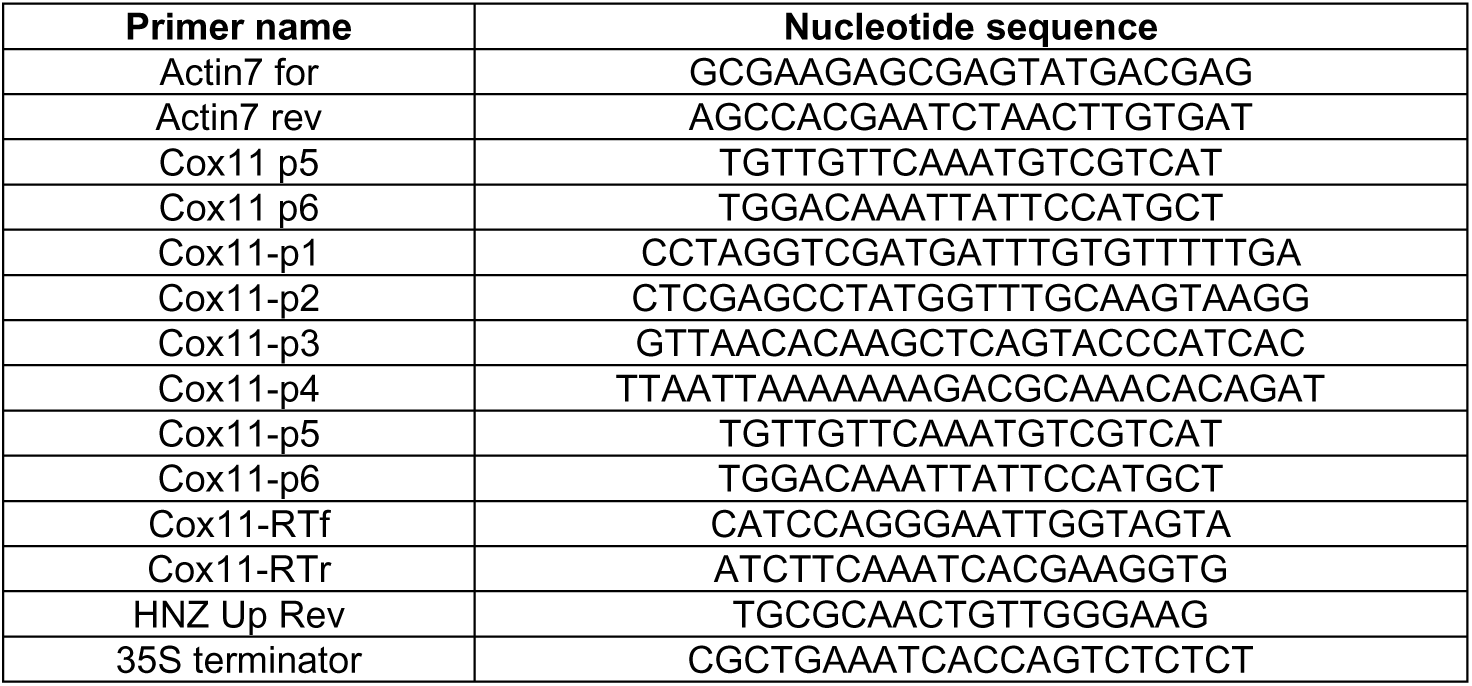
Primers used in this work. 3’→5’ nucleotide sequences are given.

**Supplementary Table 2.** Antibodies used in this work. For each antibody, the target protein complex and subunit are indicated. For commercial antibodies, the item reference is included; for homemade antibodies, the original publication is cited.

| Target complex | Target protein | Source/reference |
| --- | --- | --- |
| Complex I | Nad9 | Lamattina et al., 1993 |
| Complex II | SDH 1-1 | Peters et al., 2012 |
| Complex III | MPP alpha | Peters et al., 2012 |
| Complex V | Beta subunit | Peters et al., 2012 |
| AOX | AOX | Agrisera AS04 054 |
| PSII | Cp47 | Storti et al., 2020 |
| PSII | D2 | Storti et al., 2020 |
| PSI | PsaD | Agrisera AS09 461 |
| Flv | FlvB | Gerotto et al., 2016 |
| ATP synthase | $\gamma$ ATPase | Agrisera AS08 312 |
| PSII | PsbS | Storti et al., 2020 |
| PSI | PsaA | Agrisera AS06 172 |
| Cyt b6/f-complex | Cyt f | Agrisera AS06 119 |

**Supplementary Table 3.** Available information on ascorbate peroxidase encoding genes in *P. patens*, retrieved from three different publications, regarding gene name and predicted localization. Name in the most recent gene model version is given in the first column. Localization as described in each of the references are included: Cyt, cytosol; Chlo, chloroplast; Mito, mitochondrion; Perox, peroxisome.

| Reference | Ozyigit et al., 2016 |  | Maruta et al., 2016 |  | Wu & Wang, 2019 |  |
| --- | --- | --- | --- | --- | --- | --- |
| Gene ID | Name | Localization | Name | Localization | Name | Localitzation |
| Pp3c1_26270 | Phpat.001G104200 | Chlo/Cyt | APX3 | Perox | PpAPX3 | Perox |
| Pp3c1_40650 | Phpat.001G162800 | Chlo | APX4 | Chloro | PpAPX-S | Mito/Chloro |
| Pp3c17_7560 | Phpat.017G025400 | Chloro | - | - | PpAPX6-related | Chloro |
| Pp3c20_2050 | Phpat.020G011100 | Cyto | APX1 | Cyto | PpAPX2.1 | Cyto |
| Pp3c20_2100 | - | - | APX2 | Cyto | PpAPX2.2 | Cyto |

**Supplementary Table 4.** Relative expression of genes induced during UPR^ER^ in *P. patens*. Missing data (-) means that the gene was not detected as significantly expressed in any condition. (***) p<0.001; (**) p<0.01; (*) p<0.1; (n.s.) p>0.1 Source: Lloyd et al., 2018.

| ID in original ref | Gene ID | ZT0<br>lfc | ZT0<br>Signif. | ZT2<br>lfc | ZT2<br>Signif. | ZT6<br>lfc | ZT6<br>Signif. |
| --- | --- | --- | --- | --- | --- | --- | --- |
| Pp1s181_3V6 | Pp3c10_17310 | 0.77 | ** | 0.75 | * | 0.69 | ** |
| Pp1s566_63V6 | Pp3c15_22880 | 0.66 | * | 0.39 | n.s. | 0.01 | n.s. |
| Pp1s368_19V6 | Pp3c20_14730 | 0.75 | ** | 0.32 | n.s. | -0.18 | n.s. |
| Pp1s298_70V6 | Pp3c4_21130 | 0.47 | n.s. | 0.51 | n.s. | 0.34 | n.s. |
| Pp1s91_238V6 | Pp3c12_8210 | 0.13 | n.s. | 0.25 | n.s. | 0.23 | n.s. |
| Pp1s64_46V6 | Pp3c5_8550 | -0.50 | n.s. | 0.19 | n.s. | 0.23 | n.s. |
| Pp1s213_66V6 | Pp3c9_3330 | 0.14 | n.s. | 0.19 | n.s. | 0.26 | n.s. |
| Pp1s34_189V6 | Pp3c14_20430 | 0.13 | n.s. | -0.15 | n.s. | -0.03 | n.s. |
| Pp1s241_31V6 | Pp3c20_22800 | -0.41 | * | 0.25 | n.s. | 0.46 | * |
| Pp1s15_112V | Pp3c16_14150 | -0.48 | n.s. | 0.31 | n.s. | 0.30 | n.s. |
| Pp1s288_23V6 | Pp3c3_25750 | - | - | - | - | - | - |
| Pp1s34_31V6 | Pp3c14_18900 | - | - | - | - | - | - |

**Supplementary Table 5.**
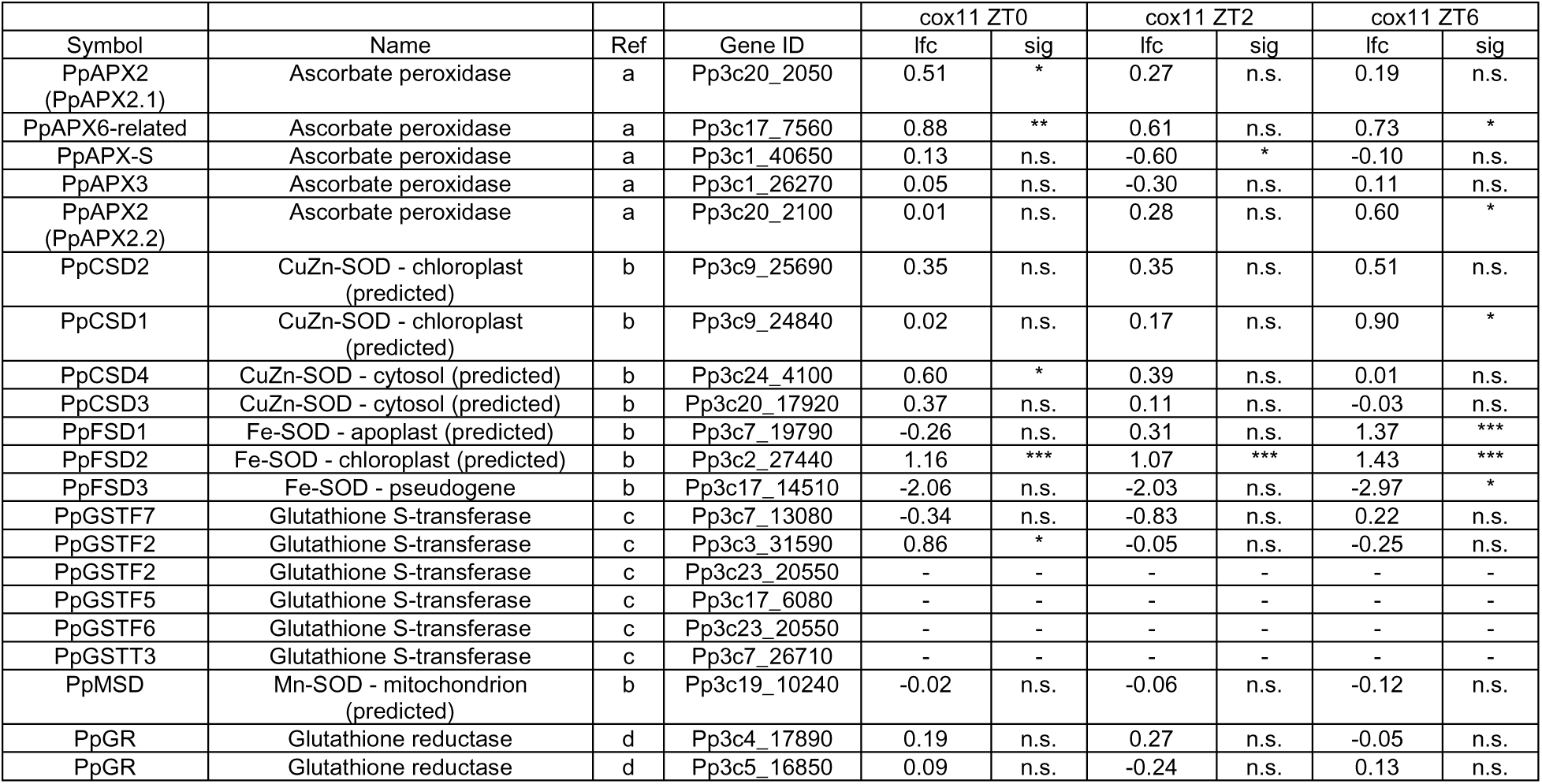
Relative expression of antioxidant enzymes in *cox11*. Missing data (-) means that the gene was not detected as significantly expressed in any condition. (***) p<0.001; (**) p<0.01; (*) p<0.1; (n.s.) p>0.1. References: (a) Wu & Wang, 2019; (b) Higashi et al., 2013; (c) Y. J. Liu et al., 2013; (d) L. Xu et al., 2013.

| Symbol | Name | Ref | Gene ID | cox11 ZT0 |  | cox11 ZT2 |  | cox11 ZT6 |  |
| --- | --- | --- | --- | --- | --- | --- | --- | --- | --- |
|  |  |  |  | lfc | sig | lfc | sig | lfc | sig |
| PpAPX2 (PpAPX2.1) | Ascorbate peroxidase | a | Pp3c20_2050 | 0.51 | * | 0.27 | n.s. | 0.19 | n.s. |
| PpAPX6-related | Ascorbate peroxidase | a | Pp3c17_7560 | 0.88 | ** | 0.61 | n.s. | 0.73 | * |
| PpAPX-S | Ascorbate peroxidase | a | Pp3c1_40650 | 0.13 | n.s. | -0.60 | * | -0.10 | n.s. |
| PpAPX3 | Ascorbate peroxidase | a | Pp3c1_26270 | 0.05 | n.s. | -0.30 | n.s. | 0.11 | n.s. |
| PpAPX2 (PpAPX2.2) | Ascorbate peroxidase | a | Pp3c20_2100 | 0.01 | n.s. | 0.28 | n.s. | 0.60 | * |
| PpCSD2 | CuZn-SOD - chloroplast (predicted) | b | Pp3c9_25690 | 0.35 | n.s. | 0.35 | n.s. | 0.51 | n.s. |
| PpCSD1 | CuZn-SOD - chloroplast (predicted) | b | Pp3c9_24840 | 0.02 | n.s. | 0.17 | n.s. | 0.90 | * |
| PpCSD4 | CuZn-SOD - cytosol (predicted) | b | Pp3c24_4100 | 0.60 | * | 0.39 | n.s. | 0.01 | n.s. |
| PpCSD3 | CuZn-SOD - cytosol (predicted) | b | Pp3c20_17920 | 0.37 | n.s. | 0.11 | n.s. | -0.03 | n.s. |
| PpFSD1 | Fe-SOD - apoplast (predicted) | b | Pp3c7_19790 | -0.26 | n.s. | 0.31 | n.s. | 1.37 | *** |
| PpFSD2 | Fe-SOD - chloroplast (predicted) | b | Pp3c2_27440 | 1.16 | *** | 1.07 | *** | 1.43 | *** |
| PpFSD3 | Fe-SOD - pseudogene | b | Pp3c17_14510 | -2.06 | n.s. | -2.03 | n.s. | -2.97 | * |
| PpGSTF7 | Glutathione S-transferase | c | Pp3c7_13080 | -0.34 | n.s. | -0.83 | n.s. | 0.22 | n.s. |
| PpGSTF2 | Glutathione S-transferase | c | Pp3c3_31590 | 0.86 | * | -0.05 | n.s. | -0.25 | n.s. |
| PpGSTF2 | Glutathione S-transferase | c | Pp3c23_20550 | - | - | - | - | - | - |
| PpGSTF5 | Glutathione S-transferase | c | Pp3c17_6080 | - | - | - | - | - | - |
| PpGSTF6 | Glutathione S-transferase | c | Pp3c23_20550 | - | - | - | - | - | - |
| PpGSTT3 | Glutathione S-transferase | c | Pp3c7_26710 | - | - | - | - | - | - |
| PpMSD | Mn-SOD - mitochondrion (predicted) | b | Pp3c19_10240 | -0.02 | n.s. | -0.06 | n.s. | -0.12 | n.s. |
| PpGR | Glutathione reductase | d | Pp3c4_17890 | 0.19 | n.s. | 0.27 | n.s. | -0.05 | n.s. |
| PpGR | Glutathione reductase | d | Pp3c5_16850 | 0.09 | n.s. | -0.24 | n.s. | 0.13 | n.s. |

**Supplementary Dataset 1 List of differentially expressed genes (DEGs) in *cox11* at different zeitgeber times**. (A) List of genes significantly regulated at each ZT. (B) List of DEGs grouped by the expression patterns described in the main text.

**Supplementary Dataset 2 List and gene composition of enriched pathways in *cox11* at different zeitgeber times.** For each pathway the following information is given: expression pattern, unique ID, ZT of regulation, source, direction, adjusted p-value, number of included genes, name of the pathway and list of included genes.

**Supplementary Dataset 3 List of metabolites identified and their relative levels in *cox11* at different zeitgeber times.** When available, IDs from multiple databases are included. Quality tags mean: A, verified with chemical standard; B, identified by database match (MS2 or MS1); C, unclear identification; D, potential degradation products (or educts).

## Acknowledgements

We thank Matteo Soldera (University of Padova) for technical assistance. Metabolite analyses were supported by the CEPLAS Plant Metabolism and Metabolomics laboratory (CMML), which is funded by the Deutsche Forschungsgemeinschaft (DFG, German Research Foundation) under Germany’s Excellence Strategy EXC-2048/1 under project ID 390686111 and CRC TRR 341 “Plant Ecological Genetics”). We also thank Elisabeth Klemp, Katrin Weber and Maria Graf (CMML) for excellent technical support.

## Author contributions

T.M. and A.A. conceived the research. A.M.V.V., A.A. and T.M. designed the experiments. A.M.V.V., M.M., L.G., P.W., A.S. and E.B. conducted the experiments, A.M.V.V., M.M., L.G., P.W., E.B., S.C., F.S., A.P.M.W., A.A. and T.M. analysed the data. A.M.V.V and T.M. wrote the manuscript.

